# Multi-layer guilt-by-association-based drug repurposing by integrating clinical knowledge on biological heterogeneous networks

**DOI:** 10.1101/2022.11.22.517225

**Authors:** Dongmin Bang, Sangsoo Lim, Sangseon Lee, Sun Kim

## Abstract

Computational drug repurposing attempts to leverage rapidly accumulating high-throughput data to discover new indications for existing drugs, often by clarifying biological mechanisms with relevant genes. Leveraging the Guilt-by-association (GBA), the principle of “similar genes share similar functions,” we introduced *clinical* neighbors of drug and disease entities while learning their mechanisms on the *biological* network. To overcome the hurdle of connecting drugs and diseases through large and dense gene-gene network and simultaneously realize the concept of “semantic multi-layer GBA”, we present a random walk-based algorithm with a novel clinical-knowledge guided teleport. As a result, drug-disease association prediction accuracy increased up to 8.7% compared to existing state-of-the-art models. In addition, exploration of the generated embedding space displays harmony between biological and clinical contexts. Through repurposing case studies for breast carcinoma and Alzheimer’s disease, we demonstrate the potential power of multi-layer GBA, a novel perspective for predicting clinical-level associations on heterogeneous biomedical networks.

## 1 Introduction

Novel drug development process in the modern era is costly, both in terms of resources and time. Drug repurposing utilizes the already-approved drugs to treat diseases, and it is increasingly becoming an attractive alternative for treatment-lacking conditions. The benefits of using ‘old’ drugs lie in the lower risk of toxicity-related clinical failure, along with lower development costs and shorter approval timelines^1^.

Accumulating bioassays and screening results have led to a more than ever understanding of drugs and diseases at the molecular level. Computational drug repurposing has gained attention owing to its rapidness and ability to utilize high-throughput data^2^, especially with the rise of the pandemic era^3^. Throughout the COVID-19 pandemic, a number of computational methodologies have been successful in finding its cures; an expert-curated network analysis discovered baricitinib^4^, which is now approved by the FDA for combination with remdesivir^5^. Furthermore, a transcriptome, proteome, and human interactome-integrative network approach along with population-based study identified melatonin as a potential prevention and treatment for COVID-19^6^. As the number of drug repurposing cases grew, so did the interest in a systematic (hypothesis-free) screen of all known drugs by fully incorporating the large bioassay datasets^7^.

Many models have attempted to connect drugs to candidate disease spaces by constructing drug-disease bipartite similarity networks^8–11^. For example, MVGCN^11^ constructs a multi-view drug-drug and disease-disease similarity network for drug-disease association (DDA) prediction. However, the limitation of these methods is that they do not fully consider the biological mode of action (MoA) of drugs and their relationship with disease.

A more convincing and widely-used method is clarifying biological mechanisms with relevant genes. This method has been well applied in the aforementioned cases of baricitinib and melatonin against COVID-19, where the target genes of the disease have already been intensively identified. However, this is not the case in general, where drug’s MoA needs to be inferred and this inferred MoA needs to be connected to disease. Hence, a *single* computational framework that connects through all three layers of drug, gene, and disease networks is required.

The systemic inference of novel DDA is mostly performed by crossing the integrated drug-target, disease-gene, and protein-protein interaction (PPI) networks. Several studies have been proposed to leverage the drug-gene-disease heterogeneous network for DDA prediction and drug repurposing^4,12–15^. Himmelstein et al.^12^ performed meta-path based network mining on a constructed heterogeneous network named Hetionet for drug repurposing. Also, Ruiz et al. analyzed the network diffusion profile of drugs on their constructed Multi-scale Interactome (MSI) network and revealed that integrating gene ontology (GO) annotations on the network improved both DDA prediction performance and interpretability. A Graph Convolutional Network (GCN)-based drug repurposing model, biFusion^16^, reported performance enhancement when the PPI network was integrated into a drug-disease bipartite network. Lastly, a recently proposed model, designated iDPath^17^, adopted a deep learning framework to connect drugs and diseases through a multi-layer biological network for drug repurposing. iDPath identified critical paths that match drugs’ MoA, implying that connection of drug and disease through the MoA-relevant path is critical for accurately predicting DDAs.

However, current biological network-based drug repurposing frameworks do not utilize the similarities between drugs and similarities among diseases in a single computational framework. Inferring functions of a biological entity through looking at its neighbors has been a consistently and widely used approach, often referred as “Guilt-by-association (GBA)”^18^. This idea of “similar entities share similar functions” is the cornerstone of biological network-based inference algorithms, including network propagation^19^. However, lifting the concept of GBA for drug repurposing is not straightforward and indeed, no existing methods are known for heterogeneous GBA that utilizes drug similarities and disease similarities.

The major difference between GBA for protein function inference and drug repurposing is that a protein and its function lie on the same layer, whereas a drug’s function is based on its *biological* level targets while their target diseases are associated at *clinical* level. To incorporate the GBA principle for drug repurposing, we aimed to realize “semantic multi-layer GBA” for drug repurposing. The core idea of multi-layer GBA is to assign the roles of a drug/disease entity by looking at its clinical neighbors, along with their topology on the biological network.

However, learning drug-disease association with the PPI network brings forth technical challenges. The main hurdle is that the PPI-based gene-gene network is much larger and denser than drug-gene and disease-gene networks. In particular, cross-network links have been reported to be highly sparse compared to the abundant PPI bioassays^20^, meanwhile the PPI network contains a sufficiently high signal-to-noise (S/N) ratio and is often too large for many algorithms^21^. Statistics of several biomedical heterogeneous networks show that gene-gene network covers over 90% of nodes and edges owing to its large number of entities and high degree (Supplementary Fig. 1). This rises the difficulty of connecting two sparse networks through a large and dense network. Owing to this limitation, network representation learning frameworks of other domains suffer from bias towards the PPI network. Our random walk and network propagation analysis on the drug-gene-disease network empirically demonstrated the algorithms’ bias towards the PPI network (Supplementary Fig. 1).

In an effort to overcome the hurdles of connecting drugs and diseases through large and dense gene-gene network and implement semantic multi-layer GBA, we applied teleport operation on drugs and diseases to populate paths passing drug and disease nodes. The original concept was introduced by the PageRank algorithm^22^, which teleports a random walker to any random node in the network. Inspired by this concept, we extend the teleport to *a semantically guided teleport* so that related drugs and related diseases can be associated. With this extension, we propose a novel framework that allows random walker to teleport to clinically similar drugs and diseases. The basis of this approach is that clinical drug neighbors share disease-relevant biological targets, demonstrated by preliminary network analysis results (Supplementary Fig. 2). This idea is identical to the core principle of translational bioinformatics: the integration of multi-scale data, as reviewed by Altman^23^. By introducing clinical neighborhood to the network, random walk paths are generated with both biological and clinical perspectives, leading to representation learning that reflects molecular and clinical contexts of entities.

Based on these ideas, we propose **DREAMwalk**: Drug Repurposing through Exploring Associations using Multi-layer random walk. DREAMwalk implements random walk with a clinical knowledge-guided teleport for heterogeneous network representation learning, and ultimately infers novel DDAs for drug repurposing.

Throughout this paper, we demonstrate the following characteristics of clinical neighborhood-based multi-layer GBA; A drug’s role can be accurately predicted with clinical neighbor information, and the generated embedding space shows harmony between clinical and molecular level contexts. Additionally, drug repurposing case studies on breast carcinoma and Alzheimer’s disease reveal the potential repurposability of candidate drugs, and ultimately, the power of multi-layered GBA principle in translating molecular world to clinical drug-disease associations.

## 2 Results

### 2.1 DREAMwalk integrates clinical information on biological network for multi-layered GBA

According to the GBA principle, characteristics of a particular biological entity can be inferred by reflecting upon its neighbors. This may be the case for single-layer GBA, such as gene or protein function prediction. However, in the case of DDA, function of a drug is defined through its biological targets and MoA; yet, its indications corresponds at a clinical level. To infer a drug’s role from both biological MoA and semantic similarity-based GBA, we introduce the concept of “semantic multi-layer GBA”.

To associate drugs and diseases, we need to generate paths across three layers; drug, gene, and disease networks. A widely used technique is to generate paths from drug to gene to disease by conducting random walks through three layers. The random walk approach is a successful method for shallow embedding of graphs. Random walk-based approaches first sample node sequences, and then pass them to representation learning architectures, for example, CBOW or Skip-gram. Owing to its flexible and stochastic nature, the algorithm demonstrates superior performance in a number of settings^24,25^. DREAMwalk fully utilizes this flexibility for integrating clinical level information and successfully implements the multi-layered GBA principle on biological networks. The overall framework of DREAMwalk is as shown in Figure 1.

**Figure 1.**
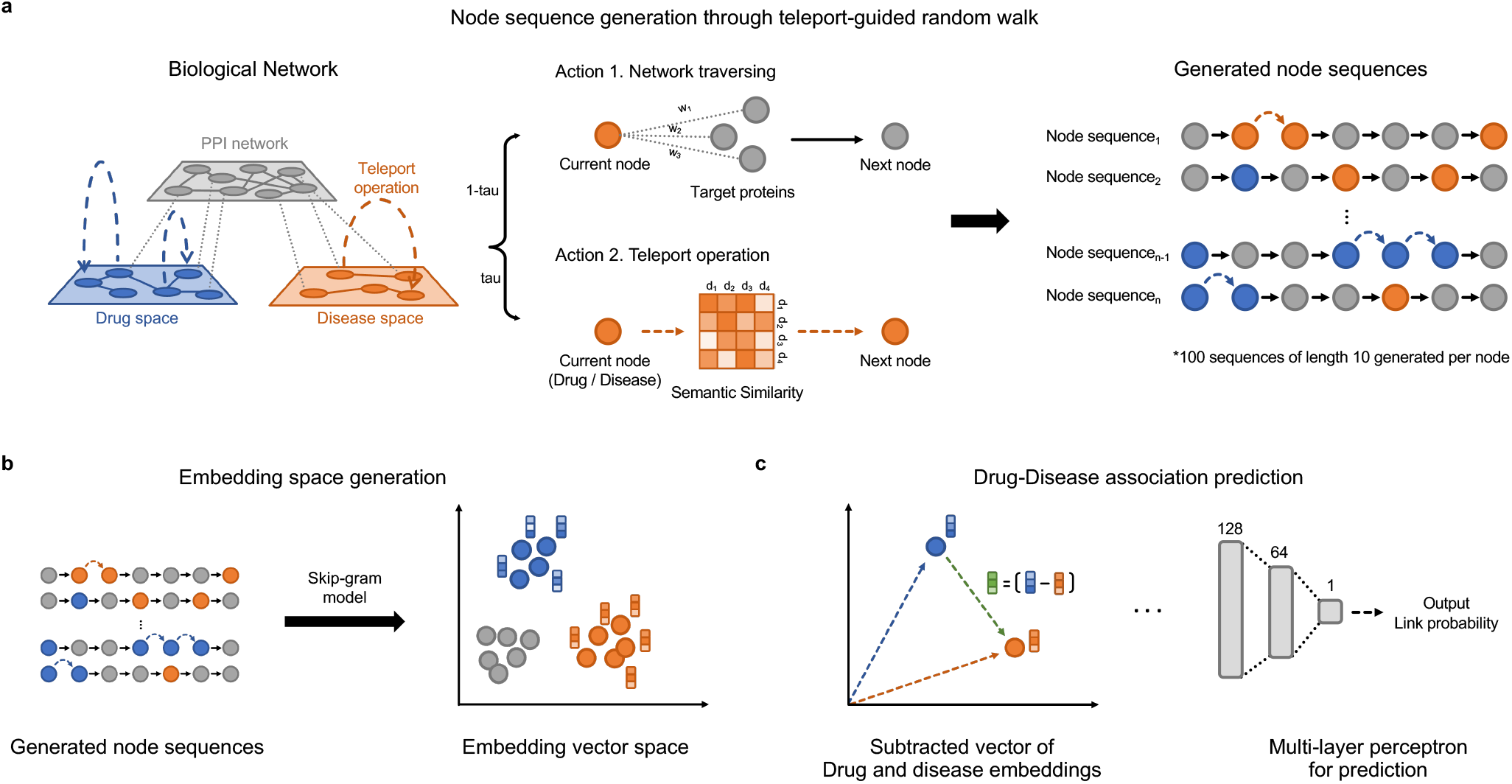
The overview of DREAMwalk framework. **a** The node sequence generation process through teleport-guided random walk. When arriving at drug/disease node, the random walker selects an action between network traversing and teleport operation based on teleport factor *τ*. **b** The embedding space generation process with Skip-gram model. **c** Drug-disease association prediction using Multi-layer perceptron with subtracted vector of drug and disease embedding vectors as input.

Inspired by the PageRank^22^ algorithm, DREAMwalk performs the random walk process with teleport operation by using clinical similarity as its guide. The widely used Anatomical Therapeutic Chemical (ATC) classification and medical subject headings (MeSH) describe the clinical hierarchy of drugs and diseases, respectively. Calculation of the semantic similarity between entities outputs the similarity matrices *S*_*drug*_ and *S*_*disease*_.

While exploring the network, when the random walker arrives at a drug or disease node, it selects its next action between *network traversing* and *teleport operation*. If network traversing action is selected, the random walker proceeds with network traversing procedure as it has done so far. If the selected action is teleport operation, the random walker randomly samples the next node from the similarity matrix *S*_*drug*_ or *S*_*disease*_, using similarity values as its sampling distribution. The probability of choosing teleport operation over network traversing is defined by the teleport factor *t*, which is a user-given parameter. This guided teleport operation leads the random walk sequence from the local neighborhood of biological level network to a clinically relevant neighborhood based on clinical similarity.

The clinical knowledge-integrated random walk sequences are then passed on to the Skip-gram model-based node representation learning. Then, using the generated node representations, a multi-layered perceptron (MLP) classifier receives the subtracted vector of drug and disease nodes and is trained to output the drug-disease treatment probability. The trained MLP model is then utilized for drug repurposing by prioritizing highly probable treatment drug-disease relationships. A detailed explanation of the model is illustrated in the Methods section.

### 2.2 Multi-layer GBA enables accurate prediction of drug-disease associations on three different biological networks

Prior to drug repurposing, we first evaluated the drug-disease association prediction performance of DREAMwalk on three biological networks: MSI, Hetionet and KEGG. In the preprocessing step, all drug-disease treatment associations were removed and left out as positive samples for the training step. An equal number of negative drug-disease pairs were randomly sampled from the network. Using the positive and negative pairs, 10-fold cross validation (CV) was performed to measure the model performance.

The selected comparison models can be clustered into random walk-based models and graph neural network (GNN)-based models, and similarity-based models. Random walk-based models consist of node2vec^26^, edge2vec^27^ and residual2vec^28^. Node2vec^26^ performs a biased random walk, whose sampling strategy can be balanced between breadth-first sampling (BFS) and depth-first sampling (DFS). Edge2vec^27^ adopts an edge-type transition matrix to consider various edge types and their semantics in heterogeneous biological networks. Residual2vec^28^ is a state-of-the-art graph representation learning algorithm proposed to debias the learning process from high-degree hub nodes by leveraging random graph sampling. Residual2vec is performed using heterogeneous node types (*hetero*-residual2vec), in addition to homogeneous node types (*homo*-residual2vec). After retrieving node sequences from each random walk-based models, the sequences are then passed on to the pipeline of Skip-gram and MLP of identical structure for drug-disease treatment association prediction.

Four GNN-based methods were used for comparison; GCN^29^, GraphSAGE^30^, graph attention network (GAT)^31^, and heterogeneous graph attention network (HAN)^32^. All models were implemented to predict the link existence between the given drug-disease pair, using PyTorchGeometric^33^.

Several models integrate multiple levels of similarities for DDA prediction and drug repurposing^8,10,11^. These methods eliminate the PPI network and construct a drug-disease bipartite network. Similarity networks include those constructed through by neighborhood similarity, i.e. Jaccard similarity of associated genes. To demonstrate the superiority of leveraging the entire PPI network, we also compared DREAMwalk’s performances with three similarity-based method: biological similarity, semantic similarity, and both similarities. The detailed description of methods and model structures for each comparison model are provided in Supplementary Methods.

The prediction performances were measured with area under receiver operating characteristic curve (AUROC) and area under precision-recall curve (AUPR). The results are shown in Figure 2. DREAMwalk outperformed state-of-the-art graph-learning methods and baseline models for the three biological networks. DREAMwalk achieved an average AUROC of 0.876 and AUPR of 0.869 on three networks, outperforming edge2vec, the best performing model in walk-based models with an average AUROC of 0.840 and AUPR of 0.839, as well as GAT, the best model among GNN-based models with average AUROC of 0.775 and AUPR of 0.767. In addition, among similarity-based approaches, the case that utilized both biological and semantic similarity networks achieved the best performance with average AUROC of 0.798 and AUPR of 0.815. Comparison of proposed model and similarity-based methods demonstrates that the necessity of the PPI network lies not only in biological interpretability, but also in performance improvement.

**Figure 2.**
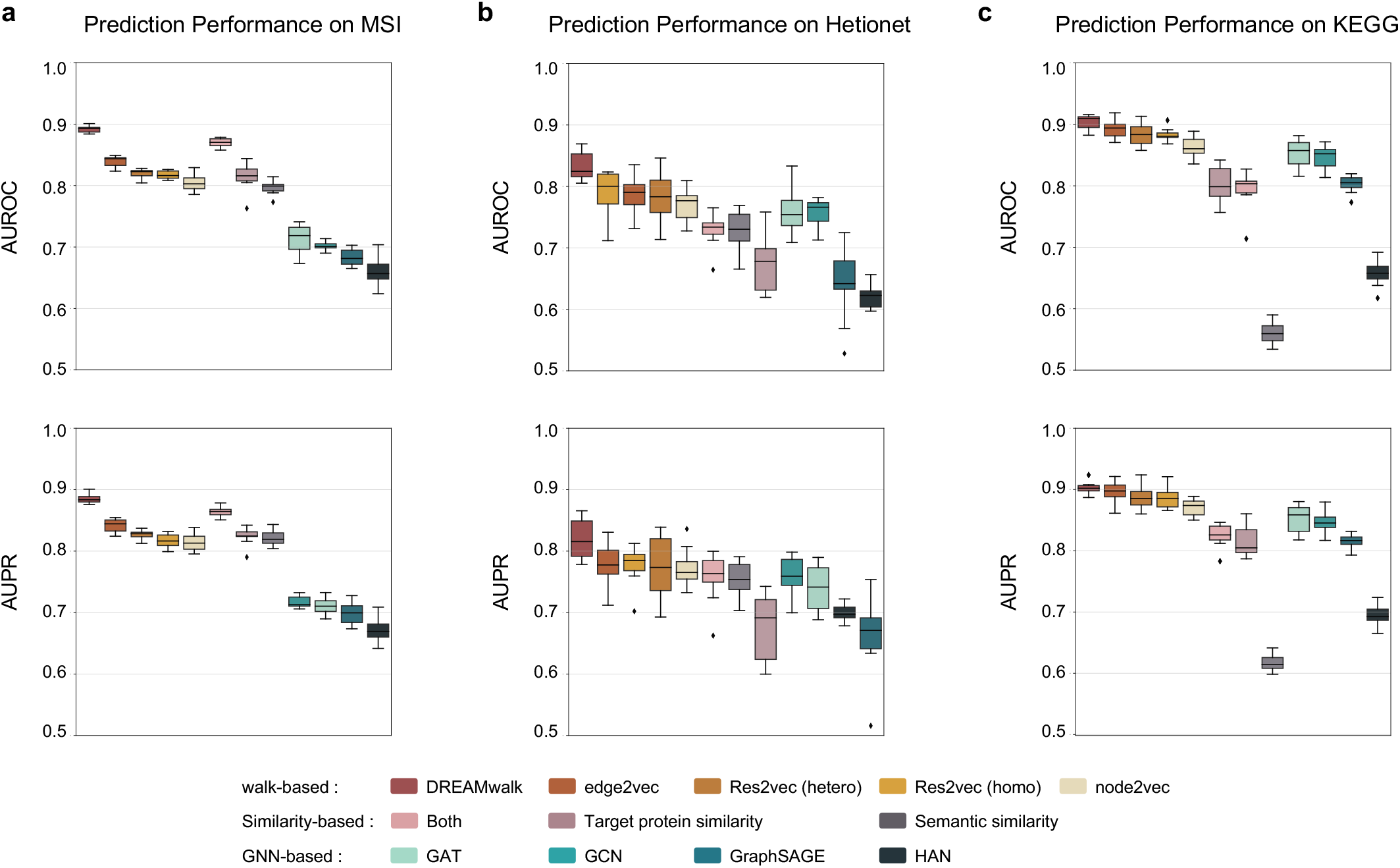
The drug-disease association prediction performances of each models on the three biological networks. AUROC: area under receiver operating characteristic curve, AUPR: area under precision-recall curve

Overall, for the three heterogeneous biological networks, DREAMwalk outperformed all other comparison link prediction models in our analysis (AUROC 0.876), ahead of state-of-the-art methods such as edge2vec (0.840), residual2vec (0.830), and HAN (0.646) and similarity-based approaches (0.815). The integration of clinical-level semantic information with teleport operation showed accurate and consistent prediction of DDA, along with its generalizability shown in three biological networks.

### 2.3 Embedding space of DREAMwalk exhibit the harmony between biological and clinical level information

We further investigated the embedding space generated by DREAMWalk to evaluate its represention of the harmonious characteristics of biological -and clinical-level information. All results reported in this section are those of the MSI network.

Investigations with multiple perspectives were performed to evaluate the embedding space of DREAMwalk, implemented *with* teleport operation, by comparing it with embedding space that is constructed *without* teleport operation. We first observed the capability of DREAMwalk’s generated space in distinguishing drug nodes from disease nodes, and also drugs by their pharmacological classes (Supplementary Fig. 3).

For a more detailed investigation of the generated embedding space, two case studies were performed to identify the characteristics of multi-layer GBA at different levels; pharmacological and systemic pathway levels.

#### Case study 1

The first case study was performed on the pharmacological level and investigated hypertensive drugs of three classes: calcium channel-blockers (CCBs), *β* -blockers (*β* Bs) and diuretics. Amlodipine, labetalol and furosemide were chosen as representative drugs of each class. CCBs, including amlodipine, lower blood pressure through inhibiting calcium channels on the surface of vascular smooth muscle cells, leading to vasodilation^34^. Labetalol, as well as other *β* Bs, treat hypertension by directly acting on the *β* -adrenergic receptors of the heart and reducing its stress^35^. Finally, diuretics, including furosemide, inhibit the reabsorption of ions and water in the kidney, resulting in increased diuresis and decreased blood volume^36^. As mentioned above, these three drug classes have different MoAs, hence they target different proteins, perturb different pathways and result in varying cellular events. The three drugs exhibits no interactions between their target proteins (Figure 3d). However, they share the same disease target: hypertension, are among the first-line treatments, and are often used in combinations^37^. These characteristics of drugs with same target disease-different MoAs may be a hurdle for biological network-based drug repurposing.

**Figure 3.**
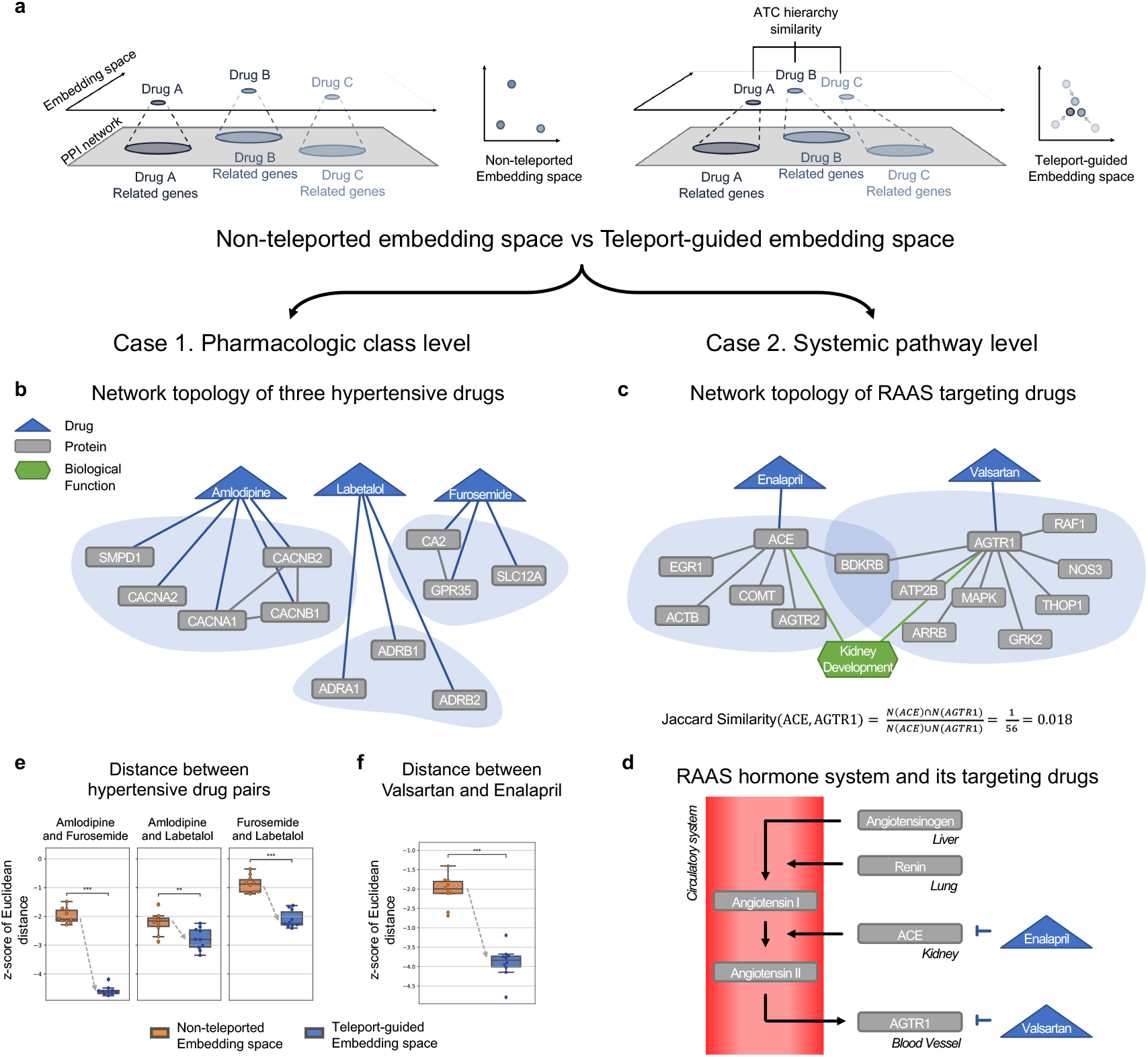
Characteristics of multi-layer Guilt-by-association shown in embedding space. **a**,**b** Network topology of the three hypertensive drug classes on without-teleport embedding space (top) and DREAMwalk embedding space (bottom). **c**,**d** Network of RAAS and its two targeting drugs. **e** The normalized euclidean distance between the hypertensive drug pairs on DREAMwalk embedding space (blue) and without-teleport embedding space (orange). **f** The normalized euclidean distance between the RAAS targeting drugs. The distance between the drug pairs are significantly decreased with the integration of clinical level annotations.

We hypothesized that in the DREAMwalk’s embedding space, the three drugs would be located close to each other since their clinical roles are analogous, even though biological MoAs differ (Figure 3c). To validate our hypothesis, normalized Euclidean distances between the three drugs were measured on both DREAMwalk’s space (generated with teleport) and space generated without teleport. Each space was generated for 10 times with different random seeds because the random walk algorithm is stochastic. As shown in Figure 3g, the measured distances of three pairs display significant reduction with the integration of clinical-level information. Since the three drugs with different MoAs but same disease targets are located closer in the multi-layer embedding space, the GBA principle can be applied in a more reasonable way for drug repurposing.

#### Case study 2

The next case study was performed at the systemic pathway level. Enalapril and valsartan are drugs that target proteins in a same hormone system, known as renin-angiotensin-aldosterone system (RAAS) (Figure 3f). Enalapril is an angiotensin-converting enzyme inhibitor (ACEI), and valsartan belongs to the pharmacological group of angiotensin receptor blockers (ARBs). Since the MoAs of the two drugs exist in the same system, they both treat hypertension, and are clinically recommended to be used separately, as their combination is associated with more adverse effects without offering any increase in benefits^38,39^. However, on the biological network, they do not share targets. In addition, the Jaccard similarity of the PPI neighbor set of ACE and AGTR1 was 0.018 (Figure 3e), implying a notable biological distance between the two drugs.

The measured normalized Euclidean distance of the two drugs significantly decreased with the integration of clinical-level information (Figure 3h). Their *cellular* pathways are well implied in the embedding space constructed without clinical teleport since the networks already contain pathway or molecular function entities. However, *systemic scale* pathway interactions, for example, hormone systems, do not appear to be sufficiently contained in biological networks. Hence, our case study demonstrates the practicality of the multi-layer GBA approach in narrowing down this gap between molecular and systemic levels.

The results of these case studies suggest that clinical knowledge-guided multi-layer GBA embedding space of DREAMwalk captures information at different scales.

### 2.4 Clinical prior knowledge-guided teleport is essential for performance improvement in DREAMwalk

DREAMwalk’s multi-layer GBA strategy, which integrates clinical knowledge into biological networks, is implemented though a novel teleport-guided random walk algorithm. To demonstrate the efficiency and significance of knowledge-guided teleport operation in accurate DDA prediction, we conducted several ablation studies.

Integrating clinical information, such as clinical hierarchies of drugs (ATC classification) and disease (MeSH term, Disease Ontology, ICD-11) is a key principle of the DREAMwalk framework. To compare the model performances with equal amount of information, we integrated clinical hierarchies as nodes on the biological networks. Figure 4a illustrates the network learning with hierarchy entity nodes attached, in comparison with teleport operation.

**Figure 4.**
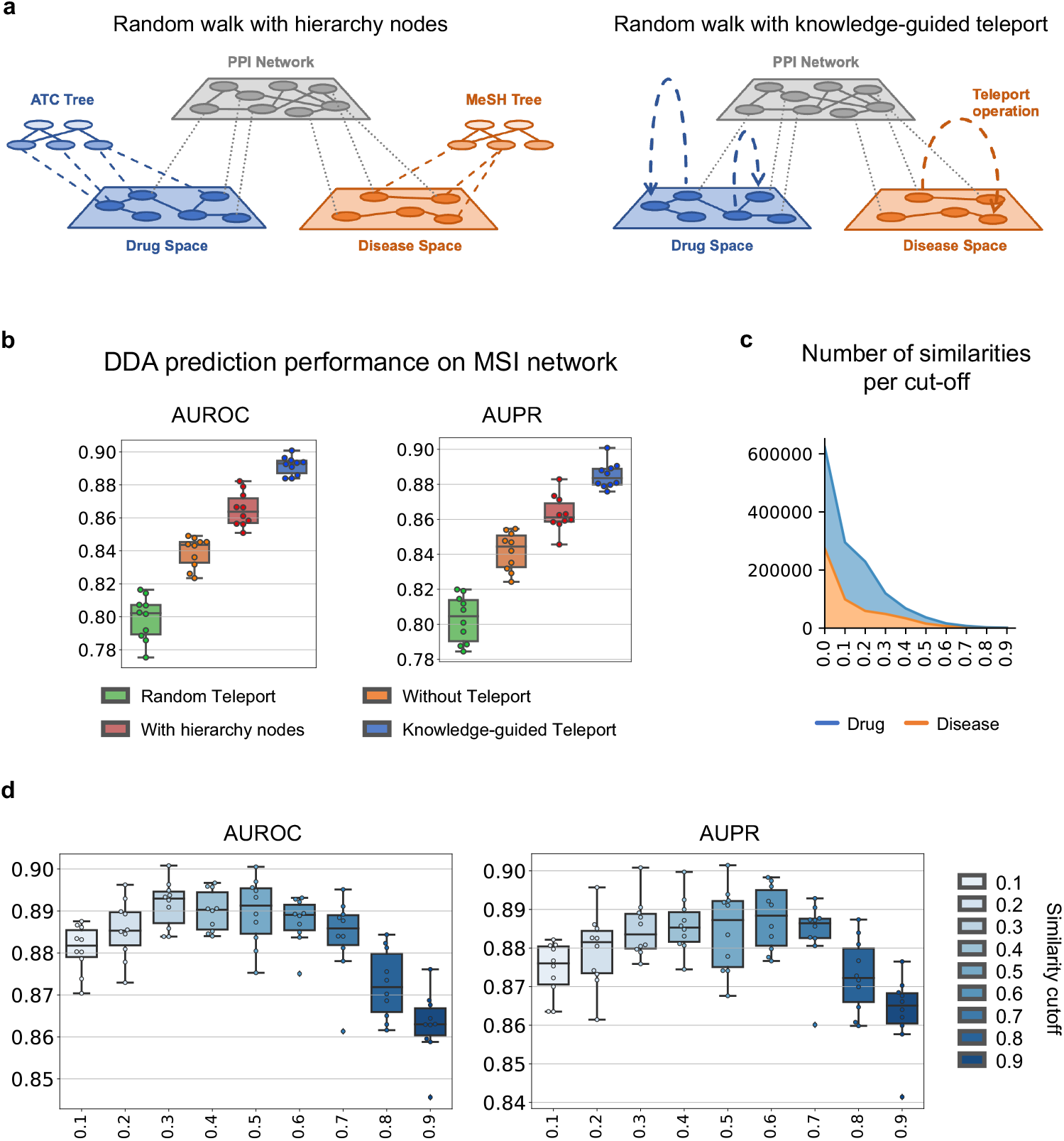
Ablation study results of DREAMwalk’s teleport operation. **a** Concept illustration of Random walk with hierarchy nodes (left) and Random walk with knowledge-guided Teleport (right). **b** Drug-disease association prediction performances of models random teleport (green), without teleport (orange), with hierarchy nodes (red) and knowledge-guided teleport (blue) on MSI network. **c** Stacked area plot of number of similarities of drug (blue) and disease (orange) per cut-off. As the cut-off value increases, the number of similarity edges used for teleport operation decreases exponentially. **d** Link prediction performances following the change in similarity cut-off on MSI network.

First, we compared the DDA prediction performances of three models: model without teleport, model with hierarchy as nodes, and model with knowledge-guided teleport on the MSI network. The results show that the integration of hierarchical information as nodes indeed results in a more accurate prediction, since additional level information is applied (Figure 4b). Notably, the increase in AUPR and AUROC was significantly higher with knowledge-guided teleport (AUROC 0.892, AUPR 0.885) compared with using the prior knowledge alone as network components (AUROC 0.865, AUPR 0.863). The performances of other baseline models on hierarchy-appended network are provided in Supplementary Figure 4.

There are advantages of integrating clinical-level information through teleport operation over learning them as network components. Along with computational efficiency due to the reduced node and edge counts (Supplementary Table 1), the signal-to-noise (S/N) ratio may be controlled by applying a cut-off value to the similarity matrix. Introducing the whole hierarchy as network component possesses the limitations of uncontrollable S/N ratio. By applying cut-off or threshold value for similarity matrix construction, the number of links are reduced (Figure 4c), leading to not only narrowing of search space but also removal of noise from irrelevant neighbors. The performances of DREAMwalk model based on different similarity cut-off values are shown in Figure 4d. The performance increased as cut-off increased until 0.6, implying a decrease in S/N ratio as dissimilar entities are excluded from teleport-able neighbors. Also, teleport factor *t* can be modified for balancing representation learning between biological and clinical levels and controlling S/N ratio (Supplementary Fig. 5).The capability of DREAMwalk algorithm in setting the cut-off point to the right value and maintaining the S/N ratio makes it more powerful tool in accurately predicting DDAs.

Another study was performed to investigate the contribution of clinical knowledge towards performance enhancements. As previously mentioned, the biological networks are heavily biased to the PPI network, since the number of nodes and their degree are much higher than the other components (Supplementary Figure 1). We hypothesized that the teleport operation’s nature of linking one drug node to another and one disease node to another may have contributed the most to the improved performance by debiasing the network learning process from PPI, instead of the clinical knowledge implied within. To determine whether the use of clinical prior knowledge contributed to the increase in performance, an additional experiment is conducted by performing teleport operation randomly. When random teleport-guided random walker selects its action as teleport, it selects the next node from randomly generated transition weights instead of clinical similarity matrix as transition weights. The random generation of transition matrix is performed for 10 times, and its performances compared with models without teleport and with knowledge-guided teleport are provided in Figure 4b.

The result was quite surprising, as we hypothesized the teleport operation’s debiasing of the learning process from PPI may contribute to improved performance. However, the performance of random teleport model (AUROC 0.799, AUPR 0.803) was significantly lower than that of the without-teleport model (AUROC 0.839, AUPR 0.842), let alone the knowledge-guided teleport model (AUROC 0.892, AUPR 0.885). Furthermore, the learning process showed instability, as the variation of the performances of 10 models are reported higher (AUROC 1.259%) than without-teleport (AUROC 0.877%) and knowledge-guided teleport (AUROC 0.528%) models. This leads to the conclusion that a clinically relevant guide is necessary for teleport operation to exert its potentials, and using both adequately results in synergistic improvement in drug-disease association prediction.

### 2.5 DREAMwalk’s clinical semantics-enriched path generation enables interpretation of drug/disease mechanisms

Leveraging biological networks for learning and prediction of drug-disease associations offers interpretability, compared to alternative ‘black-box’ learning methods. The node sequences generated during DREAMwalk’s network learning process can be analyzed to identify neighboring genes for a given entity. For analysis of neighboring genes, we defined a set of genes within a window of given size *l* in the generated node sequences (Figure 5a) of a given entity as ‘window neighbors’.

**Figure 5.**
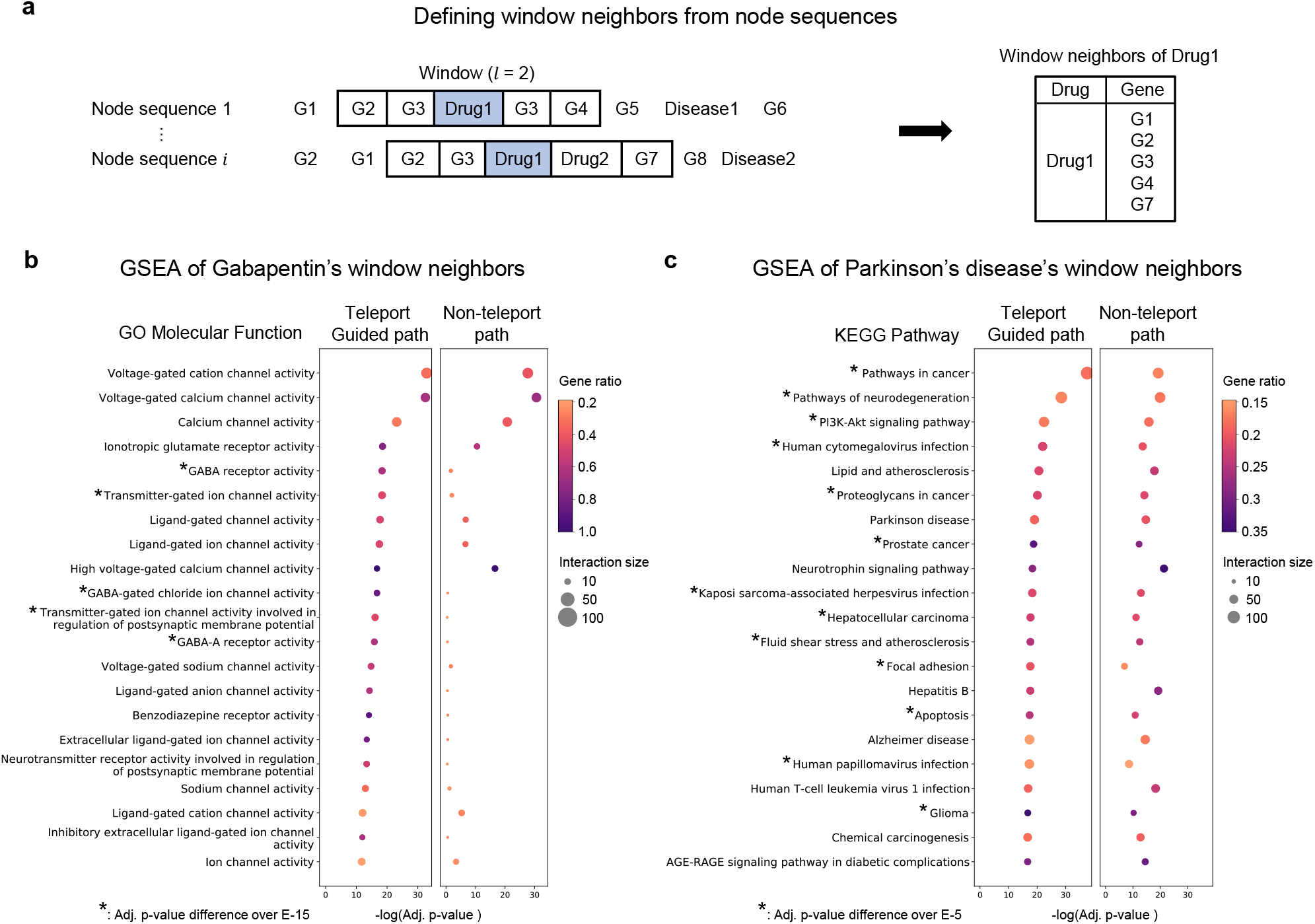
The window neighbor gene set analysis results. **a** Selection of Drug1’s window neighbor genes from node sequences using window of length 2. **b** Gene Set Enrichment Analysis results of window neighbors of drug ‘gabapentin’. **c** Gene Set Enrichment Analysis results of window neighbors of disease ‘Parkinson’s disease’. Adj.: Adjusted

Teleport-guided random walk of DREAMwalk is expected to not only explore the local neighborhood of the PPI network but also broaden the search range to clinically relevant regions. We first observed that teleport introduces more diversity to window neighbors compared with non-teleported paths (Supplementary Fig. 6).

Gene set enrichment analysis (GSEA) was further performed to demonstrate the biological interpretability of teleport-guided neighborhoods in explaining drug and disease mechanisms compared to non-teleported neighbors. Case studies were investigated with drug ‘gabapentin’ and disease ‘Parkinson’s disease (PD)’. Window neighbors were selected from window of size *l* = 2. The enrichment analysis were performed using Enrichr^40^.

Gabapentin is a relatively novel drug used in the treatment of epilepsy. The effects of gabapentin on brain amino acid neurotransmitters, including the major inhibitor gamma-aminobutyric acid (GABA), have yet to be elucidated. Studies have reported that gabapentin significantly increases GABA levels in the brain^41,42^. Interestingly, even though gabapentin alters and structurally mimics GABA, the drug does not seem to directly affect GABA-specific enzymes, GABA receptors, or GABA uptake^43^. Drug target databases reflect that gabapentin does not directly bind to GABA receptors^44,45^.

GO Molecular Function (MF)-enrichment was performed to examine the window neighboring genes of gabapentin on both teleport-guided and non-teleported paths. The resulting top 20 MFs based on adjusted p-values are shown in Figure 5b. As mentioned, gabapentin does not directly target GABA receptors, so GABA-related proteins are not located close to gabapentin in the biological network. Because non-teleport neighbor set is generated based on local neighbors of gabapentin, its GSEA results do not contain GABA-related MFs in the high ranks. In contrast, teleport-guided neighbor set captures GABA-related MFs. Enriched MFs that appeared only in the top 20 MFs of the teleport neighbors included GABA receptor activity (adj. p = 4.30E-19), GABA-gated chloride ion channel activity (adj. p = 1.92E-17), and GABA-A receptor activity (adj. p = 1.39E-16). Gabapentin’s GABAergic activities, although not directly encoded in drug-target interactions, are well captured through the clinical knowledge-integrated embedding space of DREAMwalk.

Parkinson’s disease (PD), one of the most common neurodegenerative diseases in the elderly, mainly occurs due to depletion of neurotransmitter dopamine. Various biological models have been proposed to explain the depletion of dopamine which leads to distinctive motor symptoms of PD. For the analysis of PD, KEGG pathway-enrichment was performed using its window neighbor set (Figure 5c). Among the top 20 enriched KEGG pathways in both neighbor sets, the pathways that appears on DREAMwalk were Apoptosis pathway (adj. p = 2.36E-19), Fluid shear stress and atherosclerosis pathway (adj. p = 2.18E-18) and Focal adhesion (adj. p = 4.10E-18). Literature validation confirmed that these pathways are closely related to PD.

Apoptosis is regarded as one of the main mechanisms of neuronal death in PD^46^. Although the specific processes of PD are not completely understood, it has been observed that these convergent mechanisms result in neuronal death through apoptosis, leading to PD’s motor symptoms^47,48^. Fluid shear stress and atherosclerosis pathway, along with lipid and atherosclerosis pathway (adj. p = 2.27E-21) represent the association between PD and atherosclerosis. Several studies have supported the association between atherosclerosis and PD, as well as other neurodegenerative disorders^49,50^. A large-scale Atherosclerosis Risk in Communities study (ARIC)^51^-based analysis reported decreased heart rate variability, a well-known cause of fluid shear stress and atherosclerosis^52^, was associated with an increased risk of PD^53^. Finally, Focal adhesion pathway is known to be associated with PD because adhesion playes a role in neuroprotection^54^ and the structure and function of the synapses^55^. A genome-wide association studies (GWAS) and gene expression-based integrative studies have also reported Focal adhesion as a consensus disease pathway in PD^56^.

The case studies of gabapentin and PD show the multi-layer GBA-guided neighborhood’s potential ability to explain biological mechanisms of drugs and diseases, which are difficult to identify solely via molecular-level neighborhoods.

### 2.6 DREAMwalk suggests potential repurposable drugs for Alzhiemers’ Disease and Breast Cancer

As our goal is to suggest the repurposing use of existing drugs, repurposing candidate drugs were selected for breast carcinoma and Alzheimer’s disease (AD) on the MSI network. For each disease, drug-disease association probabilities for all 1,661 drugs were calculated 10 times using DREAMwalk. After calculating the average probabilities, top 10 high-probable drugs were selected as candidates for drug repurposing. The top 10 drugs and their average probabilities with standard deviations (SD) are listed in Table 1, along with their original indications and repurposing evidence.

**Table 1.** Drug repurposing candidates of DREAMwalk for breast carcinoma and Alzheimer’s disease. The drug-disease association probabilities were measured 10 times, and the average and the SD of the predicted probabilities are provided in the table. Avg. prob.: average probability; SD: Standard Deviation; ALL: Acute Lymphoblastic Leukemia; CML: Chronic Myeloid Leukemia; NSCLC: Non-Small Cell Lung Cancer; NHL: non-Hodgkin lymphoma; SCLC: Small Cell Lung Cancer

| Breast Carcinoma |  |  |  |  |  |
| --- | --- | --- | --- | --- | --- |
| Rank | Drug | Original Indication | Avg. prob. | SD | Evidences |
| 1 | Hydroxyurea | CML, cancer of head and neck, sickle cell anemia | 0.9868 | 0.028 | 57–60 |
| 2 | Irinotecan | Colorectal cancer, SCLC, NSCLC | 0.9854 | 0.021 | 61–63 |
| 3 | Carmustine | Brain tumors, multiple myeloma, Hodgkin disease, NHL | 0.9851 | 0.026 | 64,65 |
| 4 | Clofarabine | ALL | 0.9817 | 0.022 | 66,67 |
| 7 | Etoposide | Germ cell tumors, Kaposi sarcoma, SCLC | 0.9777 | 0.038 | 62,65 |
| 9 | Vinblastine | Hodgkin disease, Lymphoma, NHL | 0.9722 | 0.037 | 62,65 |
| 10 | Erlotinib | NSCLC, Pancreatic cancer | 0.9711 | 0.069 | 68–70 |

**Table 1. Drug repurposing candidates of DREAMwalk for breast carcinoma and Alzheimer’s disease.**
| Alzheimer’s disease |  |  |  |  |  |
| --- | --- | --- | --- | --- | --- |
| Rank | Drug | Original Indication | Avg. prob. | SD | Evidences |
| 1 | Melatonin | Blind vision, sleep disorders | 0.9953 | 0.006 | 71,72 |
| 3 | Amantadine | Extrapyramidal disorders, Parkinson’s disease | 0.9926 | 0.016 | 73,74 |
| 4 | Piribedil | Dizziness, Parkinson’s disease | 0.9887 | 0.018 | 75–77 |
| 7 | Pramipexole | Parkinson’s disease, restless legs syndrome | 0.9822 | 0.027 | 78–80 |
| 9 | Phenibut | Anxiety | 0.9809 | 0.042 | 81,82 |
| 10 | Fluoxetine | Bipolar disorder, Depressive disorder | 0.9799 | 0.036 | 83,84 |

Top 10 candidate drugs for breast carcinoma mostly include chemotherapeutic agents that are often used off-label for metastatic breast cancers, as well as other metastatic cancers in clinic. In contrast, erlotinib is a drug for targeted therapy of non-small cell lung cancer (NSCLC) patients with EGFR gene mutation^85^. An interesting finding is that even though breast carcinoma is not associated with EGFR in the MSI network, DREAMwalk ranked erlotinib as candidate drug with a very high probability of 0.9711. An intensive review by Masuda et al.^68^ supports this prediction by reporting that approximately half of cases of triple-negative breast cancer (TNBC) overexpress *EGFR*, and targeting the protein leads to the rewiring of apoptotic signaling networks which enhances the chemosensitivity of cancer cells. An in-vitro study by Bao et al.^69^ and a case report by Singh et al.^70^ also supports the prediction of erlotinib’s repurposability for breast carcinoma.

AD is one of the most common cause of dementia, a neurodegenerative disorder of cerebral cortex and limbic system that results in mild cognitive decline and memory loss. Due to its close relationship with PD, several PD treatments are often used for AD treatment. Reflecting these characteristics, PD treatments amantadine, piribedil, and pramipexole are included in the top 10 repurposing candidate drugs for AD. An atypical case of phenibut, also known as fenigam, is interesting becuase it does not share any target protein with AD nor does it have an assigned ATC classification code. Phenibut was originally a neuropsychotropic drug first synthesized in and prescribed since the 1960s in Russia but has not been approved in the US and most other European countries^81^. Phenibut is known to target *GABBR1* and *GABBR2*, and yet these proteins are not associatEd with AD or any other approved AD drugs. However, phenibut is known for its cognitive enhancement effects, which possesses the potential to treat cognitive impairment symptoms^82^ of AD. The repurposing case of phenibut, a non-FDA approved and no ATC code-mapped drug, demonstrates that DREAMwalk has the potential to predict adequate indications for a novel drug with known protein target yet has no ATC code assigned.

In summary, literature-based evaluation from in vitro experiments to clinical case reports and off-label uses demonstrated the potential repurposability of the candidates for breast carcinoma and AD. Overall, the presented results and case studies demonstrate the usefulness of DREAMwalk in deriving new hypotheses for DDAs that would facilitate experimental and clinical validation and ultimately provide novel treatment strategies for treatment-poor diseases.

## 3 Discussion

The DREAMwalk framework implements a semantic multi-layer GBA for accurate DDA prediction and drug repurposing by introducing the clinical neighbors of drug and disease entities. By integrating clinical knowledge-guided teleport technique with the random walk algorithm, our representation learning process incorporates both molecular- and clinical-level information and generates a harmonized embedding space. The high DDA prediction performance on the three heterogeneous networks of MSI, Hetionet and KEGG demonstrates the generalizability of teleport-mediated integration of clinical and biological information. Ablation studies support this concept by demonstrating that knowledge-guided teleport is essential for prediction performance enhancement. Clinical prior knowledge injected through semantic similarity measure provides the largest performance enhancement, whereas randomly performed teleport results in poor performance.

Two case studies on the generated embedding space demonstrate that the characteristics of various biological levels are well projected within clinical neighborhood-guided multi-layer GBA. The GSEA analysis of generated path exploration with gabapentin and PD also displayed the interpretability of DREAMwalk in determining the mode of actions of drug and disease. Finally, DREAMwalk’s predicted repurposing candidate drugs for breast carcinoma and AD are well supported in literatures. There are however some potential limitations to DREAMwalk’s current DDA prediction and drug repurposing framework.

Although teleport operation offers efficient integration of clinical information into the biological network representation learning process, the teleport probability *t* is a user-specified hyperparameter and *t* is fixed to a preset value throughout the entire network. Adaptation of teleport probability based on local network topology may offer more flexible integration of clinical data. In addition, downstream DDA prediction task of DREAMwalk is trained based on randomly sampled negative drug-disease pair, owing to the lack of public data on negative drug-disease pairs, we plan to develop an adequate positive-unlabeled learning framework for more accurate DDA prediction in future studies.

In summary, our results indicate that the novel multi-layer GBA principle can be adopted for computational drug repurposing, inferring from clinical neighbors via random walk with clinical knowledge-guided teleport. We believe that our work is a demonstration of how to efficiently leverage clinical prior knowledge in machine learning frameworks on biological domain, as adequate integration of different levels of information is key to translating molecular information to clinical world. Also, our work may provide clues for pharmaceutical scientists to discover effective treatments for diseases that are currently without treatment options

## 4 Methods

### 4.1 Biological Heterogeneous Networks

Three biomedical heterogeneous networks, such as MSI^13^, Hetionet^12^ and KEGG^86^, were used for evaluating the drug-disease link prediction performances of the proposed and baseline models. Each network consists of varying types of nodes and edges. Node types other than drug, disease, gene and pathway (or biological function) were eliminated to construct a molecular-level biological network. Associations of higher level, for instance adverse effect or anatomy, were excluded during this process. Also, during node embedding generation step, all the drug-disease treatment edges are removed from the network. This allows the node representation learning process to fully incorporate the biological and semantic contexts of entities, without treatment association information. The drug-disease association pairs were latter used for downstream task of Multi-layer perceptron based DDA prediction.

#### Multi-scale interactome (MSI) network

Multi-scale interacome (MSI) network^13^ is a multi-scale heterogeneous biological network, including not only molecular-scale interaction but also their functional annotations. After constructing a multi-scaled biological network of drug, disease, protein and Gene Ontology^87^ Biological Function nodes, the authors generated diffusion profiles for each node through weighted network propagation and performed downstream analyses, e.g. drug mechanism analysis and drug-disease association prediction. The original MSI network consists of 4 node types and 4 edge-types. Since the network consists of only drug, disease, protein and GO biological function, the whole network is utilized for experiments and drug repurposing procedure of this work. The statistics and the data source of node and edge information are provided in the Supplementary Table 2.

#### Hetionet

Hetionet^12^ is a biomedical heterogeneous network of 11 types of nodes and 24 edge types, from 29 publicly available data sources. Hetionet is designed to integrate every available resource into a single interconnected data structure to assess the systematic mechanisms of drug efficacy. To use the original network to our experimental settings, node types other than drug, disease, protein (gene) and pathway were removed. The original and the processed network statistics are provided in Supplementary Table 3. It is worth mentioning the Hetionet contains a smaller number of disease nodes compared to MSI and KEGG because the disease nodes defined are at a higher or broader level. For instance, all hypertensive disorders and its relationships are summarized into single ‘hypertension’ node in Hetionet, while the MSI network contains not only ‘Hypertensive disease’ but also variations of the disorders, e.g. ‘Intracranial Hypertension’, ‘pulmonary arterial hypertension (PAH)’, ‘ocular hypertension’, and more.

#### KEGG

KEGG^86^ is the most widely-used database of expert-curated molecular- and pathway-level interaction annotations. The whole KEGG database contains 15 sub-databases of different types of entities. The systems information is contained in PATHWAY, BRITE and MODULE sub-databases, and the genomic information is contained in a latter-developed KEGG Orthology (KO) database. KEGG DISEASE database contains information of disease entities and their relationship to disease genes, carcinogens, pathogens and other environmental factors. KEGG DRUG database of approved drugs lists information of drug target information along with drug metabolism information. Of all the relationships contained in 15 sub-databases, only gene-pathway, drug-gene and disease-gene relationships were utilized. The statistics of the utilized KEGG network is provided in Supplementary Table 4.

### 4.2 Drug and Disease semantic similarities

Integrating clinical level information to biological network of drug-gene-disease enables drug-disease association prediction through multi-layer GBA perspective. To leverage the tree-structured hierarchical annotation of drug and disease nodes, DREAMwalk utilizes semantic similarity measure as teleport probability between drug-drug or disease-disease nodes. Based on public drug and disease ontologies, an information content-based semantic similarity measure was adopted for calculating the clinical similarities of drug-drug and disease-disease pairs. The detailed process for calculating the similarities are described below.

#### Drug and Disease Ontologies

The utilized drug hierarchy is Anatomical Therapeutic Chemical (ATC) classification hierarchy for all three heterogeneous networks. The ATC codes were assigned to drugs using the information provided by Drugbank^44^. Since the diseases IDs were mapped to different hierarchies in each network, two different disease hierarchies were used; Hetio’s disease entities were mapped to Disease Ontology^88^ hierarchy, MSI diseases were mapped to Medical Subject Heading (MeSH)^89^ term hierarchy, and KEGG diseases were mapped to ICD-11^90^. All the hierarchies can be regarded as a directed acyclic graph of terms.

#### Information Content

A number of measures have been proposed for calculating the similarity of entities in biomedical ontologies since the 1990s^91–93^. Some measures compare the entities’ information content (IC) when measuring their similarity. IC gives a measure of how informative an entity *c* is, based on the occurrence frequency of an entity in a given biomedical corpus, e.g. Uniprot Knowledge base^94^. More frequent an entity appears, less informative it is, so smaller IC is assigned to the entity. Calculating the IC value of an entity directly from a tree-structured hierarchy instead of a given corpus can be performed through counting *N*_*child*_, which is the number of children a term has in the hierarchy structure, as proposed by Seco et al.^95^. The IC value of a term in a hierarchy structure can be defined as following:

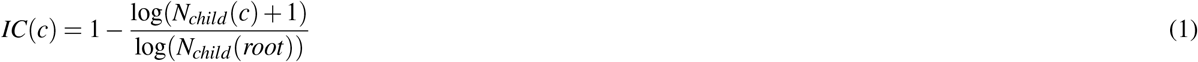

The denominator of the equation 1 assures the IC values are in [0,1], and the information content of the top entity is equal to 0.

#### Semantic Similarity

Among the most commonly used semantic similarity measures^91–93^, DREAMwalk adopted the semantic similarity measure proposed by Jiang and Conrath^91^. According to the authors, given the IC value of two entities *c*_1_, *c*_2_ and their Most Informative Common Ancestor (MICA), the distance between the two entities can be defined as following:

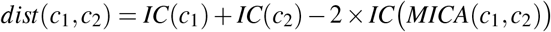

Since *max*(*IC*)= 1, the maximum value of the semantic distance between two entities is 2. In order to transform the distance measure into similarity value in range of [0,1), the similarity measure can be defined as below:

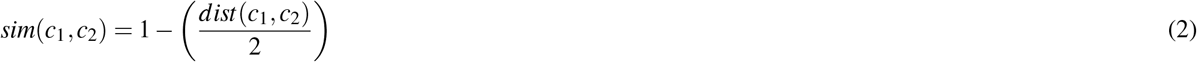

Using Equation 2, similarity measure is calculated all-pairwise for drugs and disease of the three networks, according to their drug/disease hierarchies. This procedure returns a similarity matrix *S ∈* ℝ^*n*×*n*^ where *n* is the number of drug or disease contained in a network. A user-defined cutoff may be introduced for eliminating the pair information with similarity below the given cutoff value for the reduction of noise and improvement of computational efficiency. For our study, the cutoff is empirically set to 0.3 for all networks and all similarities below are masked.

### 4.3 Multi-layered GBA through Teleport-guided random walk

Implementing the multi-layer GBA concept requires an integration of clinical level information for introducing semantic neighbors on network feature learning frameworks. To introduce clinical-level information on molecular-level heterogeneous networks, we augmented a random walk algorithm with knowledge guided teleport operation, which is inspired from the PageRank algorithm^22^.

The teleport-guided random walker generally traverses the biological network by following its edges; however, when it arrives at drug/disease nodes, it randomly selects an action between *teleport operation* and *network traversing* based on the user-given teleport factor. If the selected action is the teleport operation, the random walker teleports to a randomly sampled node based on the similarity matrix *S*. Otherwise, if the action is network traversing, the random walker resumes the traversing process. For all nodes in each network, 100 walks of length four were sampled. The detailed algorithm is provided below.

#### Random Walk

The random walk algorithm traverses nodes in the network, generating a node sequence *p* = *n*_1_, *n*_2_,…, *n*_*l*_ that can be used in the Skip-gram based graph learning framework. A node sequence of length *l* from a network *G* = (*V, E*) of node set *V* and edge set *E* can be generated by following distribution:

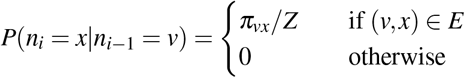

where *π*_*vx*_ is the unnormalized transition probability between nodes *v* and *x*, with *Z* as normalizing constant. The transition probability of the *un*biased random walk introduced in word2vec^96^ is equal to the edge weights *w*_*vx*_, which is equal to 1 in case of unweighted graphs.

Node2vec^26^ added search bias term *α*_*pq*_ to the transition probability which is based on parameters *p*, the return parameter, and *q*, the in-out parameter. The two parameters control the priority of the sampling strategy between breadth-first sampling (BFS) and depth-first sampling (DFS).

#### Edge-type transition matrix

To deal with the different types of edges and their semantics when generating node sequences from heterogeneous networks, an edge-type transition matrix from edge2vec^27^ is used. An edge-type transition matrix is generated based on the correlations of edge-types consisting the network through an iterative Expectation-Maximization (EM) process. Given a heterogeneous network with *m* types of edges, an edge-type transition matrix *M* ∈ ℝ^*m*×*m*^ is generated, where *M*(*i, j*) refers to the transition weight between edge-types *i* and *j*.

#### Teleport operation and teleport factor

Teleport operation is performed only when the type of current node *v, Type*(*v*) ∈ {drug, disease}. Teleport factor *t* where 0 ≤ *τ* ≤1, is a parameter that controls the rate of teleport operation and network traversing. In the proposed teleport-guided-random walk algorithm, when the random walker arrives at a drug or disease node, the teleport action is chosen with a probability of *t*; otherwise, the network traversal continues, with a probability of 1 − τ.

For instance, if τ = 0.3, the random walker on a drug/disease node has 30% probability of selecting teleport action and 70% probability of choosing network traversing. Thus, setting *t* to a high value makes teleport operation more frequent, enabling the influence of clinical similarity to become greater and vice versa.

If the selected action of the random walker is teleport operation, the next node is randomly sampled from weighted probability as calculated in the similarity matrix *S*, defined through the process described in Section 4.2. For example, given a current drug node *v*_*drug*_, the probability for the next node to be drug node *n*_*drug*_ through teleport operation can be express as below:

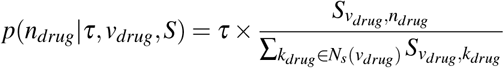

where *t* is the teleport factor, and *N*_*s*_(*v*_*drug*_) is the neighboring node set of node *v*_*drug*_ in the drug similarity matrix *S*. The same process is computed for disease nodes with disease similarity matrix *S*^*disease*^.

#### Network traversing

Network traversing is performed when: 1) the current node type is other than drug/disease, or 2) the current node type is drug/disease and the selected action is network traversing. Given the current node *v* and the previous node *u* with the trained edge-type transition matrix *M*, the probability of selecting the next node *n* can be shown as below:

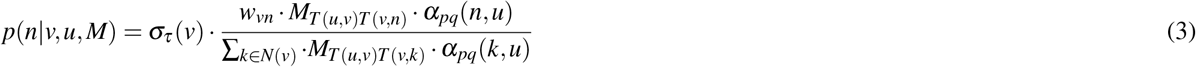

where σ_*t*_ (*v*) is the network traverse probability defined by the node type of *v, N*(*v*) is the neighboring node set of node *v* and *T* (*u, v*) is the edge-type between *u* and *v*. Network traverse probability term is defined as follows:

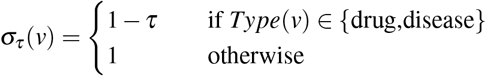

### 4.4 Node embedding generation

Skip-gram model is used for learning continuous feature representations of nodes in the generated sequences by teleport-guided random walks (Section 4.3). The representation learning of the Skip-gram model is performed by optimizing a neighborhood-preserving likelihood objective function using stochastic gradient descent (SGD) with negative sampling. The node representation steps are implemented using the Gensim^97^ python package. The `windowlength` parameter, that is the maximum distance between the current node and the predicted node in a node sequence, is set to 4 while training all models.

### 4.5 Multi-layer Perceptron Classifier for Drug-Disease association prediction

After the node representations for all biomedical entities in the network are generated, a Multi-layer perceptron (MLP) is used for drug-disease link prediction and drug repositioning.

#### Multi-layer Perceptron

MLP is a fully connected feed-forward artificial neural networks (ANN) that is widely used in various application domains including computational drug discovery^98^. For learning the relationship between drug and disease, feature vectors obtained by element-wise subtraction of embedding vectors of two entities are used as input for MLP consisting an input layer, an output layer and one hidden layer in between. The output layer consists of a single neuron wrapped by sigmoid function which passes down the predicted probability of positive drug-disease treatment association. Binary Cross Entropy (BCE) loss is used with Adam algorithm^99^ as optimizer for training procedure. Early stopping strategy is adopted with the patience of 20 epochs on the validation loss.

#### Drug-disease association prediction

The DREAMwalk framework is composed of two steps; node embedding generation step and link prediction step. As mentioned above, we removed all the drug-disease links during the first step to generate drug and disease embedding with their MoA contexts. The drug-disease pairs were used in the second step; DDA prediction task as positive set. Negative drug-disease pairs of equal number of positive pairs were randomly sampled from the network. 10-times cross validation (CV) setting was adopted, and negative sampling was conducted in a way that there are no overlapping samples between the 10 CV sets. The split ratio of train, validation and test sets was set to 7:1:2. The CV setting was applied identically to all the models evaluated for this study.

## Data availability

Each heterogeneous networks were retreieved from the corresponding GitHub repositories and through API calls; Hetionet (https://github.com/hetio/hetionet), MSI (https://github.com/snap-stanford/multiscale-interactome) KEGG (https://www.kegg.jp/kegg/rest/keggapi.html)

## Code availability

The source code for DREAMwalk’s node embedding and DDA prediction are available at the following GitHub repository (https://github.com/eugenebang/DREAMwalk).

## Acknowledgements

This research was supported by the Bio & Medical Technology Development Program of the National Research Foundation (NRF), funded by the Ministry of Science & ICT(NRF-2022M3E5F3085677) (to S.K.).

## Author contributions

D.B., S. Lim, S. Lee and S.K. conceived the experiments, D.B. conducted the experiments, D.B., S. Lim, S. Lee and S.K. analysed the results, D.B. wrote the manuscript. All authors reviewed the manuscript.

## SUPPLEMENTARY INFORMATION FOR

### Supplementary Methods 1

#### Methods for path-based comparison models

Random walk generated path-based models leverage skip-gram model for node representation learning. All models including proposed model DREAMwalk generated node embedding vectors through same skip-gram settings from sampled paths. For all models, 100 paths of length 10 were samples for each node. Latent vector dimension was set to size of 128.

##### node2vec

node2vec^1^ is a flexible random walk generated path-based method that balances between breadth-first sampling and depth-first sampling through parameters *p* and *q*. For all experiments, *p* and *q* were set to 1.

##### edge2vec

edge2vec^2^ is a node2vec-based model that considers heterogeneous edge types and performs biased path generation. Prior to biased path sampling, edge2vec first performs uniform path sampling for learning the edge type distribution of the network and generate an edge-type transition matrix. For edge-type transition matrix generation process, 1 path of length 10 were sampled for all nodes. Also, bias parameters *p* and *q* were set to 1.

##### residual2vec

residual2vec^3^ is node embedding model that leverages random graphs for debiasing networks. residual2vec’s path generation can be performed in homogeneous and heterogeneous modes. For comparison, both modes of res2vec-homo and res2vec-hetero were utilized. All parameters were set to values given by the authors.

#### Methods for similarity-based comparison models

Several drug repurposing models leverage different similarity networks for drug-disease association (DDA) prediction. For comparison, we generated target protein-similarity network and semantic similarity network and performed node2vec path sampling for node embedding generation.

#### Target similarity network

Jaccard similarity measure was utilized for generating drug-drug and disease-disease similarity networks based on their associated gene sets. Drug-drug network is generated from

#### Semantic similarity network

The measure of Jiang and Conrath^4^, modified by Seco et al.^5^, was utilized for generating semantic similarity network of drug and disease.

#### Target and semantic similarity Integrated network

Lastly, an integrated drug-disease bipartite network of both target-similarity and semantic similarity networks was constructed. Performance comparison with integrated similarity network was conducted since DREAMwalk utilizes both drug-gene-disease and semantic similarity information.

### Methods for GNN-based comparison models

Unlike path-based models, link prediction using graph neural network (GNN) models are performed in an end-to-end manner. Four GNN-based models^6-9^ were used for performance comparison. All convolution layers were stacked two times, and link prediction was performed with the inner product of drug and disease representations. All convolution layers were implemented using PytorchGeometric^10^ python package.

#### Node Features

Since GNN requires node features prior to downstream tasks, we tested the performances on three settings of node features: vector composed of four centralities (degree, betweenness, closeness and eigenvector), eigenvectors for the K smallest eigenvalues of the normalized Laplacian matrix^3^, and trainable lookup embedding layer. As a result, trainable lookup embedding layer showed the best performance and was selected for performance comparison.

#### HAN

Heterogeneous Graph Attention Network (HAN)^9^ generates node embeddings by aggregating features from meta-path based neighbors. Hence, the user needs to specify the meta-paths prior to model learning and link prediction. Since the edge types differ for three heterogeneous networks MSI, Hetionet and KEGG, we searched and found following sets of best performing meta-paths, described in Supplementary Table 5.

**Supplementary Fig. 1.**
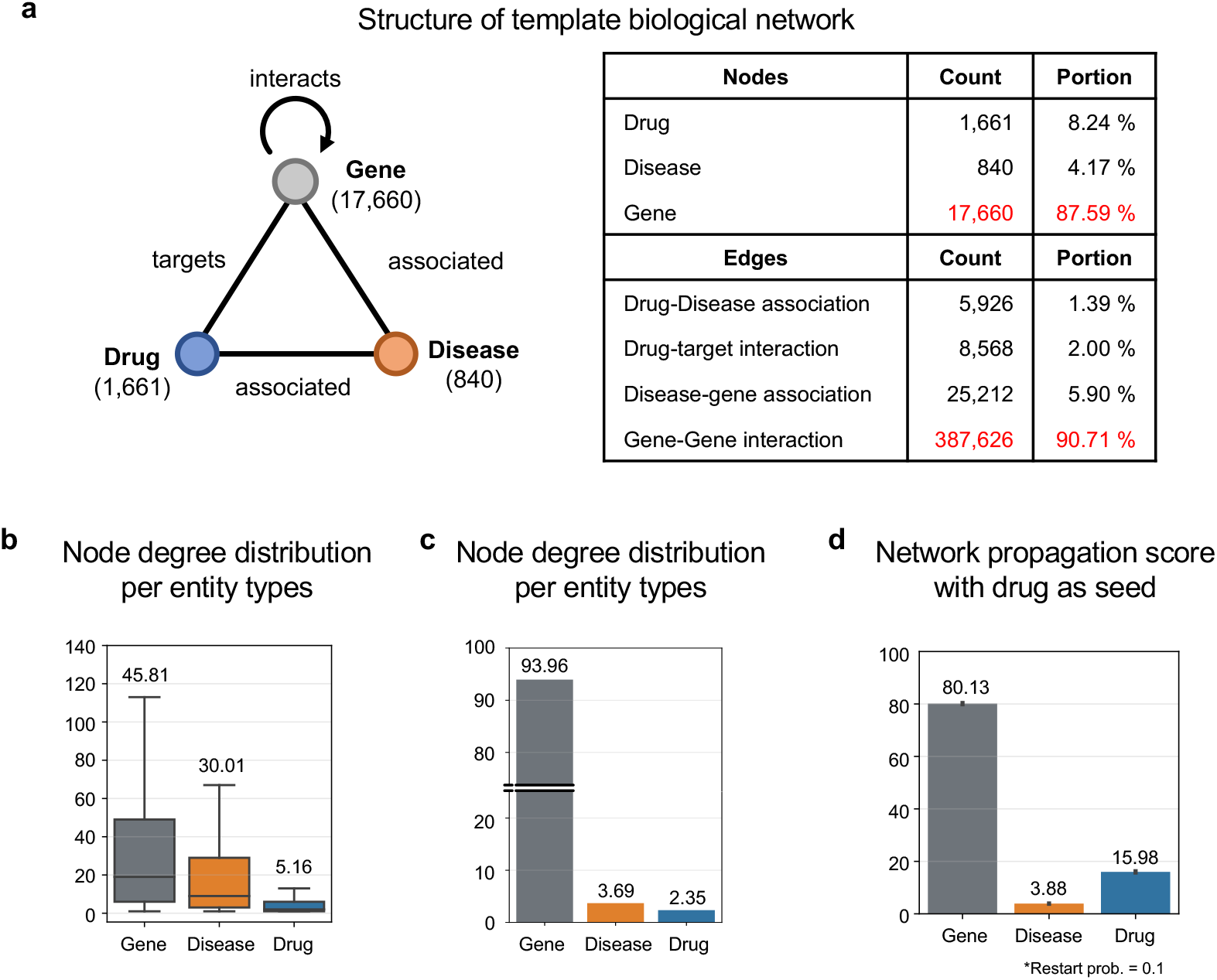
Analysis of drug-gene-disease network reveals its bias to gene-gene network. **a** Schematic and statistics of template drug-gene-disease network, extracted from the MSI^11^ network. Genes cover up over 87% of the nodes and gene-gene interaction edges cover up over 90% of the edges on the whole network. **b** Node degree distribution for each node types. The degrees of gene nodes are higher (average 45.8) than that of drugs (5.2) and diseases (30.0). **c** Node type distribution from sampled random walk sequences. 10 walks of length 10 were generated for each node uniformly. **d**. Network propagation scores for each node type, performed with Random walk with Restart (RWR) algorithm. The RWR was performed for all drug entities as seeds with restart probability of 0.1. The average score is notated above each plot.

**Supplementary Fig. 2.**
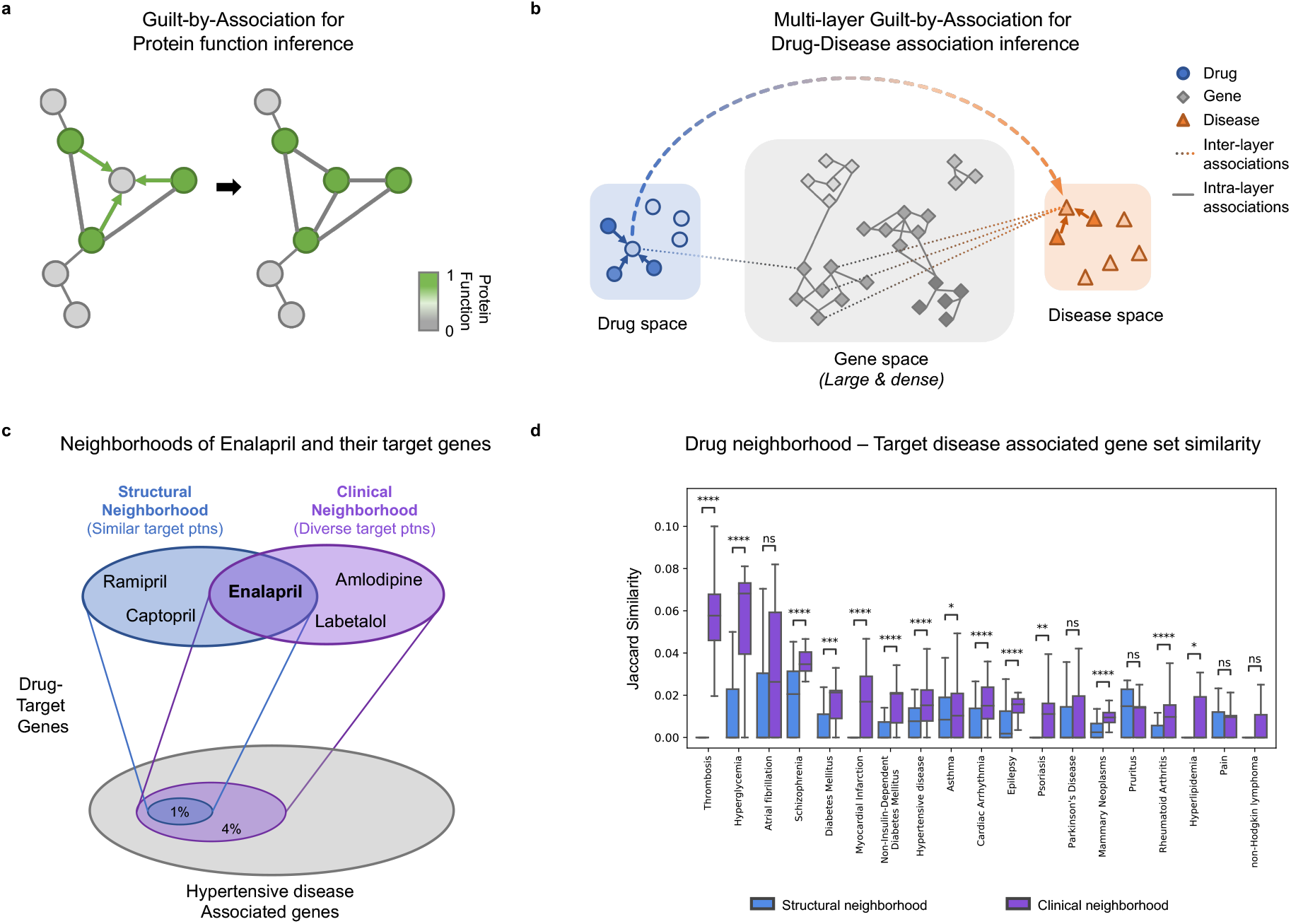
Multi-layer Guilt-by-Association (GBA) and network analysis of drug neighborhoods revealing protein targets of a clinical drug neighborhood cover a wider area of target disease-associated genes. **a** GBA for protein function inference through looking at a protein’s interaction neighbors. **b** Multi-layer GBA for drug-disease association inference through looking at a drug’s neighbors. **c** Concept of hypertensive drug enalapril’s structural neighborhood and clinical neighborhood, and their target genes relative to hypertensive disease-associated genes. **d** Gene set similarity of neighborhood target genes and disease-associated genes. Target genes of clinical neighborhood show higher similarity with target disease-associated genes compared to structural neighborhood. ns: p >= 5e-2; *: 1e-2 < p < 5e-2; **: 1e-3 < p < 1e-2; ***: 1e-4 < p < 1e-3; ****: p < 1e-4

**Supplementary Fig. 3.**
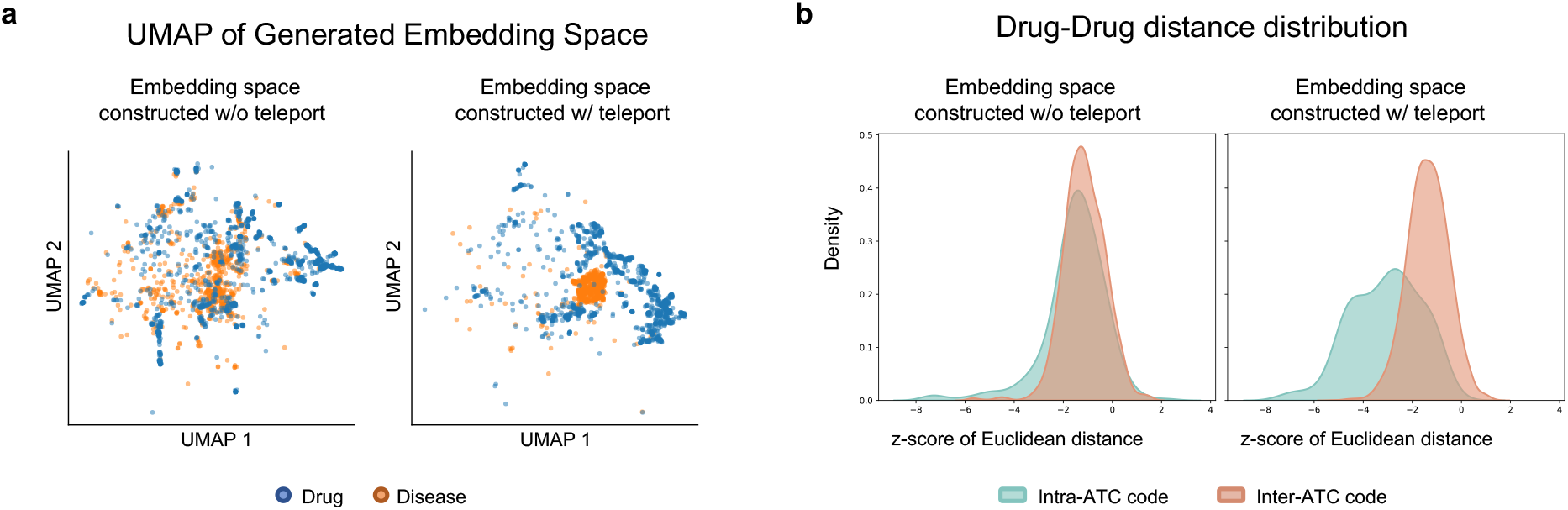
Embedding space of DREAMwalk exhibits its ability to distinguish entities based on their semantics. **a** The UMAP plot of without-teleport embedding space (left) and DREAMwalk embedding space generated with teleport (right). Drug (blue) and disease (orange) entities are well separated in the DREANwalk’s embedding space. **b** Comparison of inter-ATC level 1 distances (orange) and intra-ATC level 1 (blue) distances on DREAMwalk embedding space (left) and without-teleport embedding space (right). Intra-ATC code distance refers to the distances between the source and target drugs in the same first-level ATC code, whereas inter-ATC code distance refers to the distance between drugs of different first-level ATC codes. Since the first-level ATC code clusters drugs into 14 main anatomical or pharmacological groups, the multi-layer GBA space shows drugs sharing ATC annotations located closer to each other. In contrast, the distance distribution of the space generated without teleport implicit the biological level network contains insufficient information for clustering drugs based on their anatomical and pharmacological characteristics.

**Supplementary Fig. 4.**
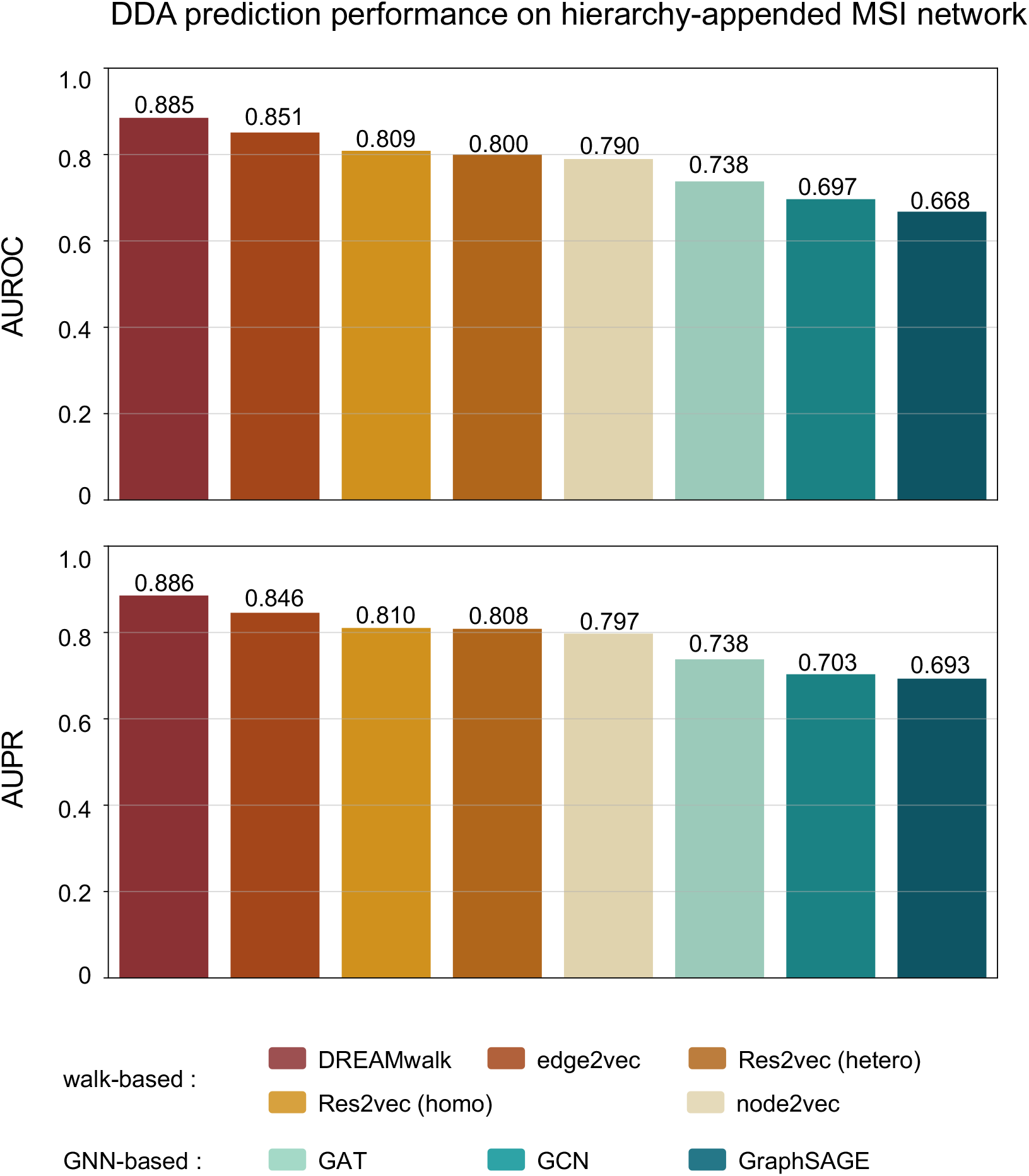
Drug-disease association prediction performances of comparison models on hierarchy-appended MSI network.

**Supplementary Fig. 5.**
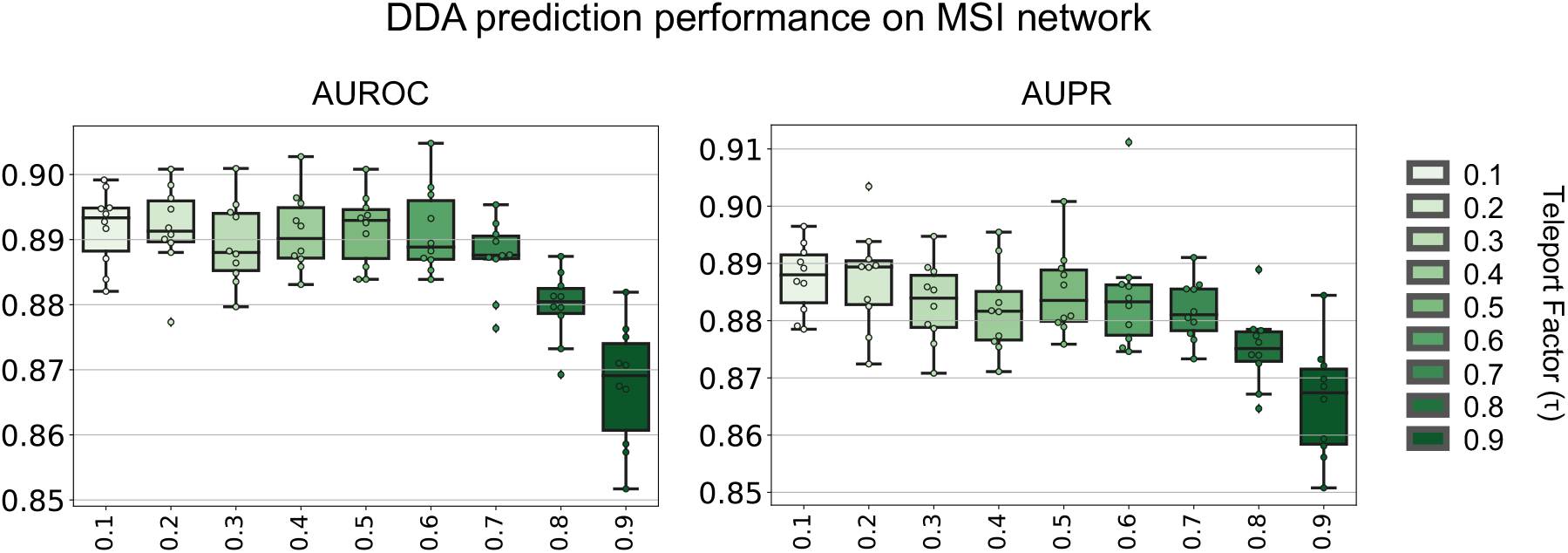
Drug-disease association prediction performances following the change in teleport factor τ on MSI network. The precise selection of teleport factors for balancing representation learning between biological and clinical levels is critical for better prediction of DDA. To find an appropriate combination of biological and clinical information, experiments were conducted several times while changing the teleport factor, τ. AUROC peaks when τ = 0.6 and drops drastically as τ increases. This suggests that applying teleport at an appropriate level contributes to enhanced performance. Too frequent or too little teleport operation leads to a performance decrease, meaning that the balance between biological and clinical data is critical when implementing their integration. DDA: drug-disease association, AUROC: Area under the receiver operating characteristic curve, AUPR: Area under the precision–recall curve.

**Supplementary Fig. 6.**
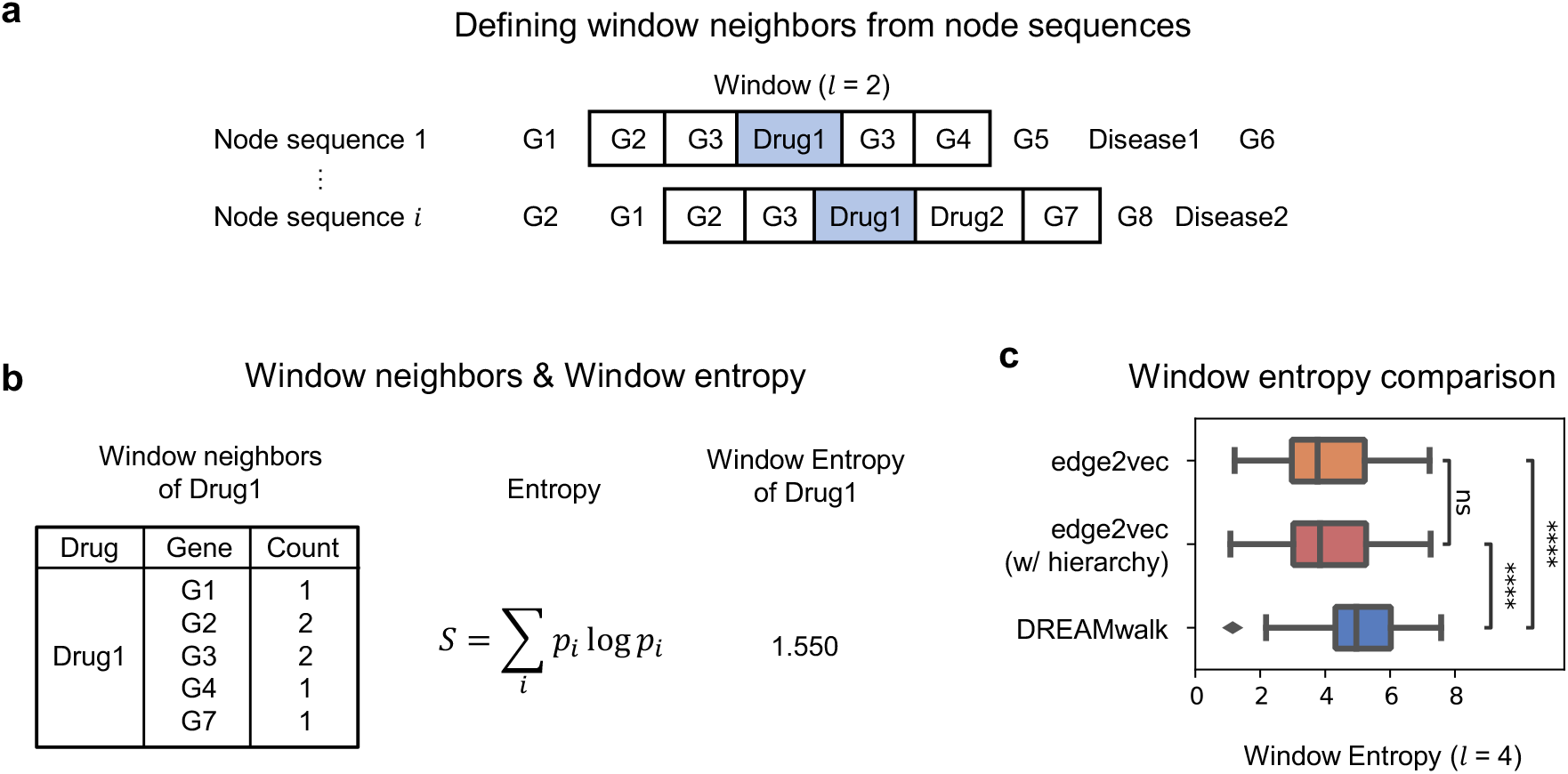
Window neighbor entropy comparison of edge2vec and DREAMwalk. **a** Selection of from node sequences with window length of 2. **b** Calculating window entropy from generated node sequences based on each neighbor’s counts. **c** Comparison of window entropy for all drug entities with node sequences generated by edge2vec (on original MSI network), edge2vec w/ hierarchy (on MSI network with hierarchy entities as nodes), and DREAMwalk. The window length was set to 4 for window entropy comparison

**Supplementary Table 1.** Statistics of utilized drug/disease hierarchies. ATC classification, MeSH term, DO and ICD-11 hierarchies for each of the three heterogeneous networks.

| Network | Hierarchy | Number of terms | Number of associations | Total levels |
| --- | --- | --- | --- | --- |
| MSI | ATC | 4,647 | 4,543 | 5 |
|  | MeSH | 3,328 | 4,373 | 12 |
| Hetionet | ATC | 3,289 | 3,274 | 5 |
|  | DO | 308 | 302 | 12 |
| KEGG | ATC | 5,314 | 5,210 | 5 |
|  | ICD-11 | 1,126 | 972 | 7 |

**Supplementary Table 2.** Statistics of the MSI network. All the nodes and edges of the original MSI network are utilized for experiments.

| Nodes |  |
| --- | --- |
| Drugs | 1,661 |
| Diseases (Indications) | 840 |
| Genes | 17,660 |
| GO Molecular Functions | 9,798 |
| Edges |  |
| Drug-disease associations | 5,926 |
| Drug-target interactions | 8,568 |
| Disease-gene associations | 25,212 |
| Protein-protein interactions | 387,626 |
| Gene-GO Molecular Function annotations | 34,777 |
| GO Molecular Function associations | 22,545 |

**Supplementary Table 3.**
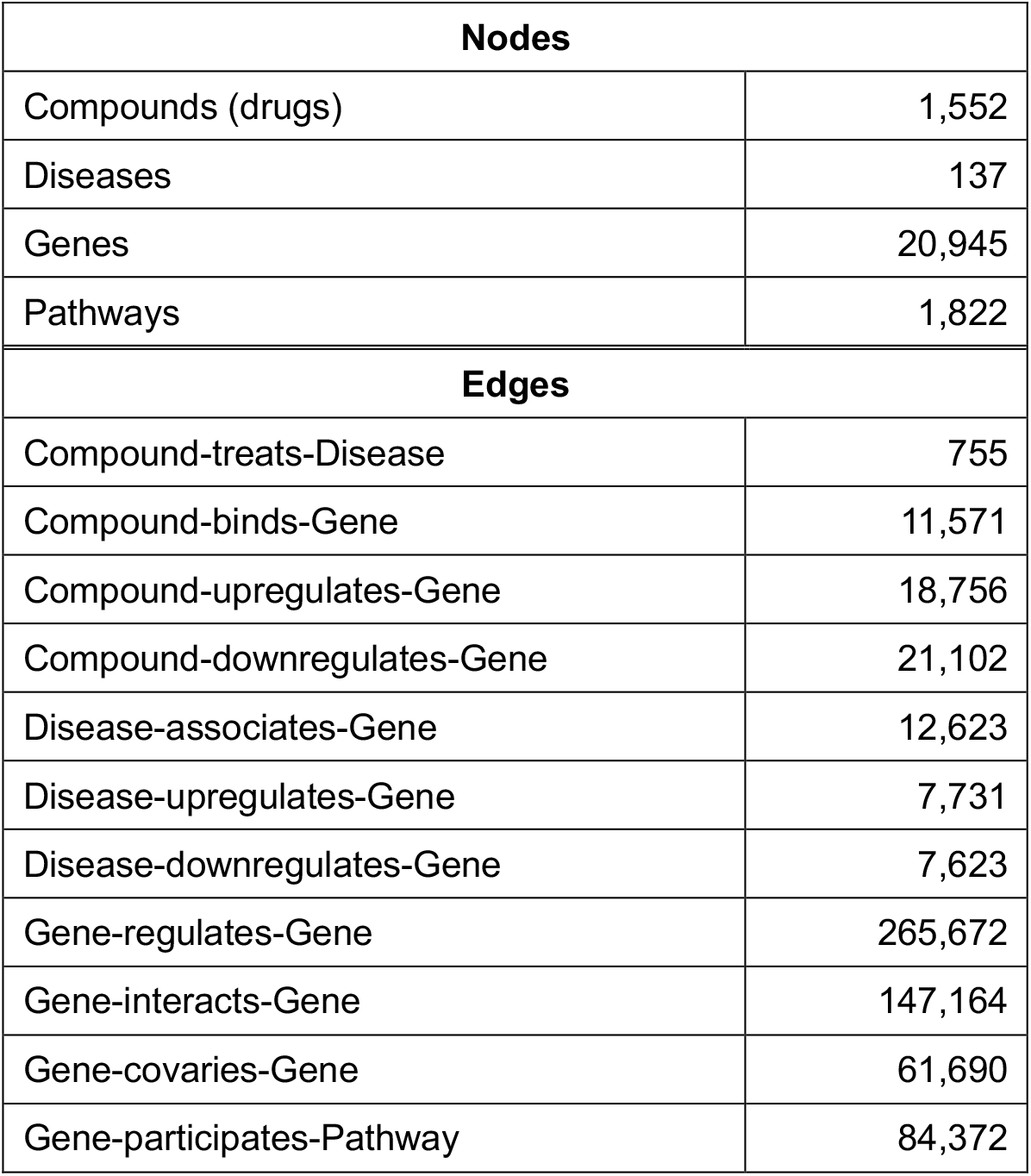
Statistics of the Hetionet network. Nodes and edges associated with drugs, diseases, genes were extracted from the original Hetionet^12^ to construct a biological level network.

| Nodes |  |
| --- | --- |
| Compounds (drugs) | 1,552 |
| Diseases | 137 |
| Genes | 20,945 |
| Pathways | 1,822 |
| Edges |  |
| Compound-treats-Disease | 755 |
| Compound-binds-Gene | 11,571 |
| Compound-upregulates-Gene | 18,756 |
| Compound-downregulates-Gene | 21,102 |
| Disease-associates-Gene | 12,623 |
| Disease-upregulates-Gene | 7,731 |
| Disease-downregulates-Gene | 7,623 |
| Gene-regulates-Gene | 265,672 |
| Gene-interacts-Gene | 147,164 |
| Gene-covaries-Gene | 61,690 |
| Gene-participates-Pathway | 84,372 |

**Supplementary Table 4.**
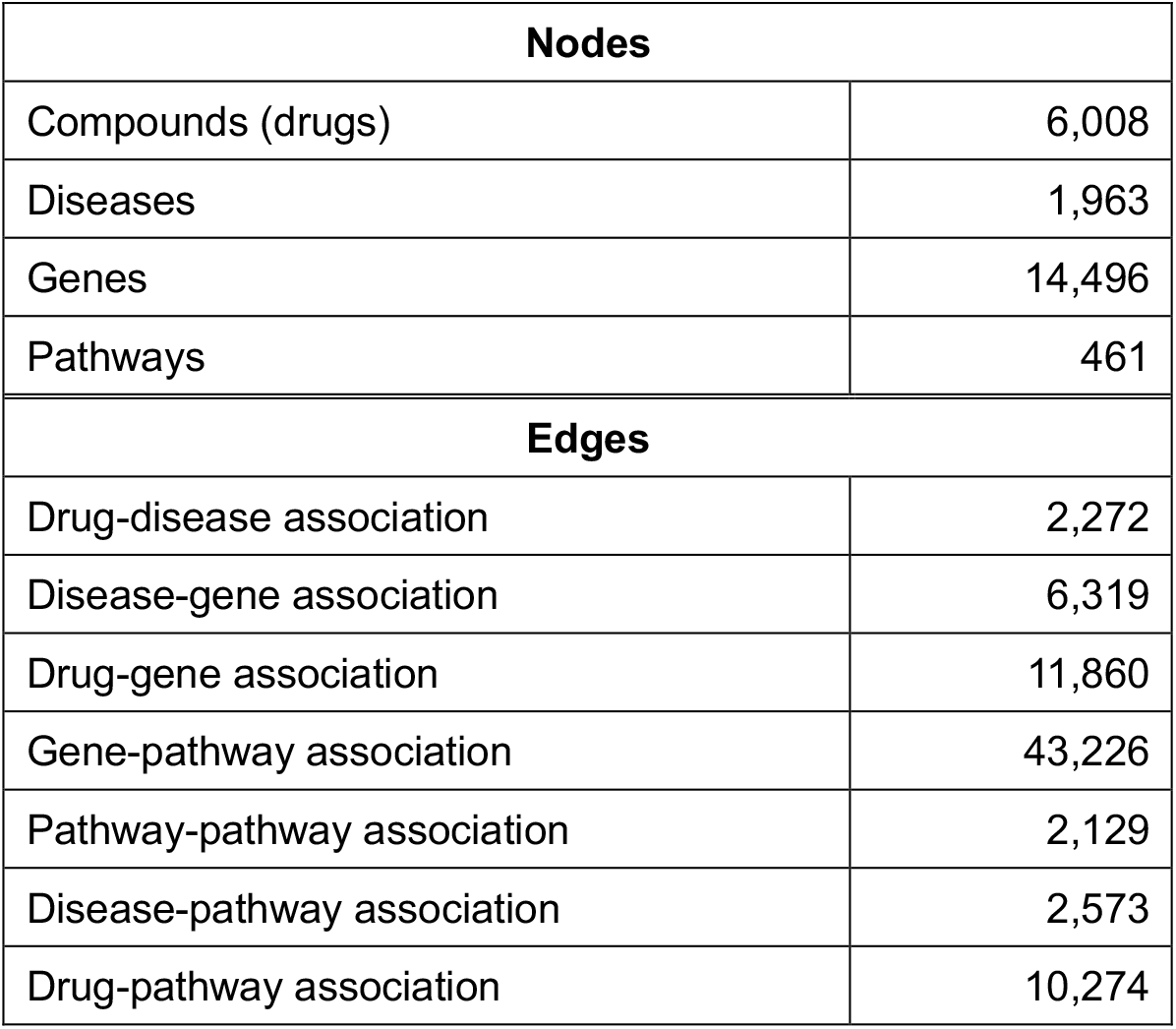
Statistics of the KEGG network. Nodes and edges associated with drugs, diseases, genes were extracted from the KEGG^13^ network.

| Nodes |  |
| --- | --- |
| Compounds (drugs) | 6,008 |
| Diseases | 1,963 |
| Genes | 14,496 |
| Pathways | 461 |
| Edges |  |
| Drug-disease association | 2,272 |
| Disease-gene association | 6,319 |
| Drug-gene association | 11,860 |
| Gene-pathway association | 43,226 |
| Pathway-pathway association | 2,129 |
| Disease-pathway association | 2,573 |
| Drug-pathway association | 10,274 |

**Supplementary Table 5.**
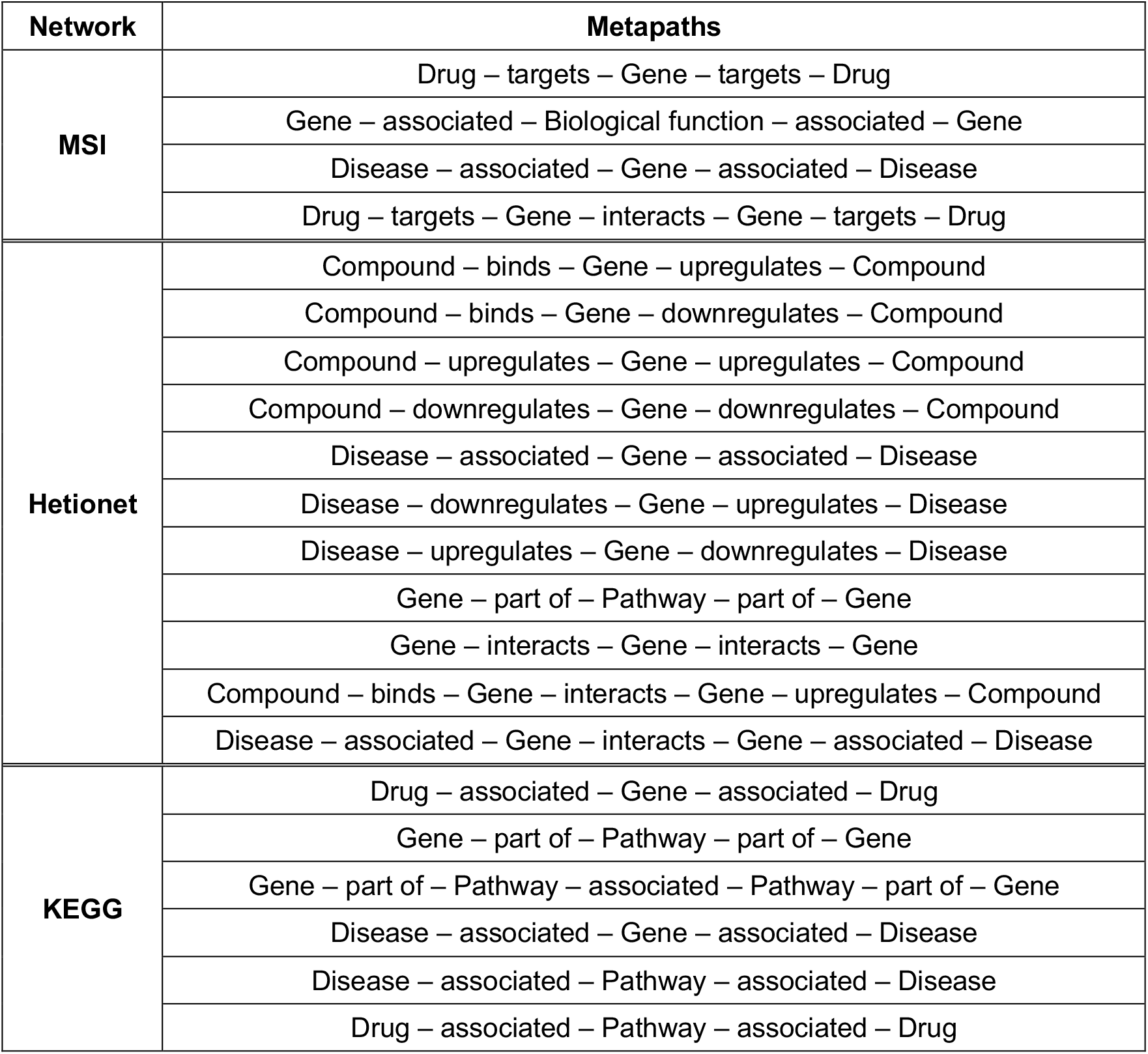
Input metapaths for HAN on each network. Since the edge types differ for three heterogeneous networks MSI, Hetionet and KEGG, we searched and found following sets of best performing meta-paths within length 3 for measuring the drug-disease association prediction performances.

| Network | Metapaths |
| --- | --- |
| MSI | Drug – targets – Gene – targets – Drug |
|  | Gene – associated – Biological function – associated – Gene |
|  | Disease – associated – Gene – associated – Disease |
|  | Drug – targets – Gene – interacts – Gene – targets – Drug |
| Hetionet | Compound – binds – Gene – upregulates – Compound |
|  | Compound – binds – Gene – downregulates – Compound |
|  | Compound – upregulates – Gene – upregulates – Compound |
|  | Compound – downregulates – Gene – downregulates – Compound |
|  | Disease – associated – Gene – associated – Disease |
|  | Disease – downregulates – Gene – upregulates – Disease |
|  | Disease – upregulates – Gene – downregulates – Disease |
|  | Gene – part of – Pathway – part of – Gene |
|  | Gene – interacts – Gene – interacts – Gene |
|  | Compound – binds – Gene – interacts – Gene – upregulates – Compound |
|  | Disease – associated – Gene – interacts – Gene – associated – Disease |
| KEGG | Drug – associated – Gene – associated – Drug |
|  | Gene – part of – Pathway – part of – Gene |
|  | Gene – part of – Pathway – associated – Pathway – part of – Gene |
|  | Disease – associated – Gene – associated – Disease |
|  | Disease – associated – Pathway – associated – Disease |
|  | Drug – associated – Pathway – associated – Drug |

